# State anxiety biases estimates of uncertainty during reward learning in volatile environments

**DOI:** 10.1101/809749

**Authors:** Thomas P Hein, Lilian A Weber, Jan de Fockert, Maria Herrojo Ruiz

**Author notes:** Corresponding author: Mara Herrojo Ruiz,. Address: Department of Psychology, Goldsmiths, University of London. Lewisham Way, New Cross, London SE14 6NW (UK).

## Abstract

Previous research established that clinical anxiety impairs decision making and that high trait anxiety interferes with learning rates. Less understood are the effects of temporary anxious states on learning and decision making in healthy populations. Here we follow proposals that anxious states in healthy individuals elicit a pattern of aberrant behavioural, neural, and physiological responses comparable with those found in anxiety disorders, particularly when processing uncertainty in unstable environments. In our study, both a state anxious and a control group learned probabilistic stimulus-outcome mappings in a volatile task environment while we recorded their electrophysiological (EEG) signals. By using a hierarchical Bayesian model, we assessed the effect of state anxiety on Bayesian belief updating with a focus on uncertainty estimates. State anxiety was associated with an underestimation of environmental and informational uncertainty, and an increase in uncertainty about volatility estimates. Anxious individuals deemed their beliefs about reward contingencies to be more precise and to require less updating, ultimately leading to impaired reward-based learning. We interpret this pattern as evidence that state anxious individuals are less tolerant to informational uncertainty about the contingencies governing their environment and more uncertain about the level of stability of the world itself. Further, we tracked the neural representation of belief update signals in the trial-by-trial EEG amplitudes. In control participants, both lower-level precision-weighted prediction errors (pwPEs) about the reward outcomes and higher-level volatility-pwPEs were represented in the ERP signals with an anterior distribution. A different pattern emerged under state anxiety, where a neural representation of pwPEs was only found for updates about volatility. Expanding previous computational work on trait anxiety, our findings establish that temporary anxious states in healthy individuals impair reward-based learning in volatile environments, primarily through changes in uncertainty estimates and potentially a degradation of the neuronal representation of hierarchically-related pwPEs, considered to play a central role in current Bayesian accounts of perceptual inference and learning.

## 1. Introduction

Anxiety is characterised by excessive worry about negative possibilities (Grupe and Nitschke, 2013). It can lead to distinct difficulties when making decisions and learning about the world, as anxious individuals experience negative reactions to uncertainty, known as intolerance of uncertainty (IU; Bishop, 2007; Carleton, 2016). Recent work has established that individuals high in trait anxiety have difficulties adapting their learning rate to changes in probabilistic task environments (Browning et al., 2015; Huang et al., 2017). Less understood is how temporary states of anxiety in healthy subjects interfere with optimal learning and belief updating in the brain. Identifying the computations that subserve learning under state anxiety is important due to the prevalence of highly anxious states in most real-world environments that are filled with uncertainty (Bach et al., 2011; Bishop and Gagne, 2018). In addition, these insights could expand our understanding of the mechanisms by which anxiety biases beliefs about the world, linking to anxiety-related disorders.

Previous computational work identified three types of uncertainty during decision-making and learning: irreducible uncertainty, informational (estimation) uncertainty, and environmental uncertainty (Bland and Schaefer, 2012; de Berker et al., 2016; Yu and Dayan, 2005). Irreducible uncertainty emerges from the probabilistic relationships between responses and their outcomes, which is an inherent property of most real-world interactions. Estimation uncertainty arises from the imperfect information about those response-outcome relationships. Lastly, environmental uncertainty induced by volatility represents the possibility of change in that probabilistic environment. The latter affects learning as the individual is uncertain about her estimates as the world is changing. To reduce uncertainty, the brain is thought to appraise the inherent statistical structure of the world using probability distributions, continuously updating and inverting a hierarchical model of the sensory inputs (de Lange et al., 2018; Doya et al., 2007; Friston, 2010; Rao and Ballard, 1999). In this context, each type of uncertainty is expressed by the width (variance, or its inverse, precision) of the probability distribution of the corresponding belief (Feldman and Friston, 2010; Mathys 2011).

Examinations of belief, uncertainty, and precision estimates using Bayesian formulations in perceptual and learning tasks are increasingly used to provide mechanistic explanations for an array of neuropsychiatric conditions. Specifically, difficulties estimating precision have been suggested to explain various clinical expressions, from movement difficulties in Parkinson’s disease to features of schizophrenia and autism (Friston et al., 2016, 2013; Lawson et al., 2017, 2014; Parr et al., 2018). In the case of anxiety, altered beliefs are also theorised to play a vital role (Paulus and Stein, 2010; Paulus and Yu, 2012). As anxiety relates to worry over uncertainty, volatile task environments have been used to understand how trait anxiety affects learning, providing a mechanistic account of anxiety-related disorders (Browning et al., 2015; Huang et al., 2017). Healthy individuals are known to adapt their learning rate to volatility, with changing environments promoting a higher learning rate as new information needs to be integrated to better predict the future (Behrens et al., 2007). By contrast, high-trait anxious individuals show reduced adaptability of their learning rate to volatile environments, both in aversive (Browning et al., 2015) and reward settings (Huang et al., 2017). Moreover, they show poorer performance in decision-making tasks (de Visser et al., 2010; Miu et al., 2008).

Expanding on those findings, here we evaluated whether temporary anxious states in healthy individuals influence reward learning in a volatile environment through changes in informational and environmental uncertainty. Evidence for a link between anxiety and inaccurate estimation of uncertainty would lend support to recent theoretical accounts suggesting that difficulties learning from incomplete information and misestimations of uncertainty are crucial to understanding affective disorders (Pulcu and Browning, 2019).

Probabilistic inference has been proposed to be achieved through the sequential use of Bayes’ rule: by dynamically combining our predictions (prior beliefs) with new evidence (sensory data), and weighting each resultant prediction error (PE) according to its precision (Feldman and Friston, 2010; Friston and Kiebel, 2009; Kok and de Lange, 2015). This predictive coding scheme relies on the hierarchical flow of information between cortical regions (Bastos et al., 2012; Iglesias et al., 2013; Rao and Ballard, 1999). Predictions are transmitted down the cortical hierarchy (backward) to meet incoming ascending (forward) sensory PEs thought to arise in supragranular layers in superficial pyramidal cells (Friston and Kiebel, 2009). Beliefs are then updated by reducing PE signals across each level of the cortical hierarchy weighted according to their estimated precision (Kok and de Lange, 2015). Importantly, developments in Bayesian computational modelling allow us to estimate inter-individual differences in the trial-wise computations and expression of these precision-weighted PEs (Mathys et al., 2014, 2011).

Monkey single-cell recording and human functional magnetic resonance imaging (fMRI) studies have shown that PEs elicited by reward are encoded by phasic responses in midbrain dopamine neurons, and these signals are conveyed to the medial frontal cortex (MFC; Chew et al., 2019; Matsumoto et al., 2007; Morris et al., 2006; Zarr and Brown, 2016). Using electroencephalography (EEG), these reward learning signals can be detectable in the error related negativity (ERN), an event-related potential (ERP) triggered by overt errors around 100 ms; and the feedback ERN (fERN) that follows negative feedback around 250 ms (Holroyd et al., 2003; Montague et al., 2004; Nieuwenhuis et al., 2004; Yeung et al., 2005). Both components have been shown to originate in the posterior medial frontal cortex (pMFC, including the anterior cingulate cortex, ACC; Holroyd et al., 2003; Montague et al., 2004; Yeung et al., 2005). Relevant to our study, the fERN has been proposed to index the magnitude of prediction violation (surprise), thus reflecting a reward PE signal that can be estimated, for instance, by using reinforcement learning models (Gehring and Willoughby, 2004; Holroyd and Coles, 2008; Holroyd and Krigolson, 2007). Also, the P300 (peaking between 250 - 500 ms) of parietal topography may be sensitive to reward PEs, valence, and surprise (Hajcak et al., 2007, 2005; Polich, 2007; Wu and Zhou, 2009).

More compelling evidence linking PEs, Bayesian surprise, and belief updating to changes in ERP responses comes from studies combining computational modelling and analysis of trial-wise EEG responses (Diaconescu et al., 2017a; Jepma et al., 2016; Kolossa et al., 2015; Mars et al., 2008; Stefanics et al., 2018; Weber et al., 2019). For instance, recent EEG studies on the MMN were able to spatiotemporally dissociate lower-level precision-weighted PE (pwPE) signals, which drive updates in belief estimates (Stefanics et al., 2018), and higher-level pwPEs, driving volatility updates (Weber et al., 2019). In addition, model-based single-trial analyses of the P300 identified the earlier P3a waveform of anterior distribution as an index of belief updating, whereas Bayesian surprise was represented in the later posterior P3b component (Kolossa et al., 2015). Here we were interested in assessing the neural representation of pwPEs across different levels, including lower-level pwPEs used to update reward tendency estimates, and higher-level pwPEs used to update volatility estimates, as belief updates on these two levels differentially depend on informational and environmental uncertainty. Accordingly, we evaluated the effect of these two hierarchically-related pwPEs on brain activity by analysing trial-wise ERP responses across frontal, central, and parietal brain regions, and within a broad temporal range from 200 to 500 ms, encompassing the fERN and P300 components.

To address our questions, we examined cortical dynamics in a control and a state anxious group using EEG recordings during a reward-based learning task. Further, to link the anxiety-induced neural changes to potential alterations in uncertainty estimation, we used a generative Bayesian inference model of perception and learning, the Hierarchical Gaussian Filter (HGF) (Mathys et al., 2014, 2011). The HGF estimates individual trajectories of trial-wise belief updates governed by hierarchically related PEs based on the responses of participants. To reveal the effect of hierarchical PEs and precision weights on evoked brain responses, we used the relevant hierarchical computational quantities (pwPEs) as regressors in a general linear model (GLM) of trial-wise EEG amplitudes as done in previous studies (Diaconescu et al., 2017a, 2014; Weber et al., 2019).

## 2. Material and Methods

### 2.1 Participants

Forty-two healthy individuals (age 18-35, 28 females, mean age 27, standard error of the mean [SEM] 0.9) participated in this reward-based learning study following written informed consent. This experiment was approved by Goldsmiths University of London’s ethics review committee. Our sample size was informed by previous computational work on anxiety (Browning et al., 2015). All participants were healthy volunteers, with no past neurological or psychiatric disorders.

All participants were screened using Spielberger’s Trait Anxiety Inventory (STAI; Spielberger, 1983) which has reliably demonstrated internal consistency and convergent and discriminant validity (Barnes et al., 2002; Spielberger, 1983; Spielberger et al., 1970). Scores on this trait inventory range from low (20) to high anxiety (80). Participants were measured for their trait anxiety level (mean 46, SEM 1.5) and then split into two groups using the median value (43). The sample population range was between 34 and 68 (Low trait = 34-42, High trait = 43-68). This created a high and low trait anxiety group to then randomly draw from to create the experimental and control groups. Importantly, trait anxiety levels did not exceed the clinical level (> 70: this cutoff score represents the mean and 2 SD above the mean for adults (Spielberger, 1983; Taylor et al., 2005).

### 2.2 Experimental Design

Using a between-subject experimental design with state anxiety being the between-subject factor, we allocated participants pseudo-randomly (after screening trait anxiety scores) to the experimental (state anxiety, StA) or control (Cont) group. They completed our experimental task, which consisted of four blocks, resting state 1 (R1: baseline), reward-learning task block 1 (TB1), reward-learning task block 2 (TB2), and resting state 2 (R2; see **Supplementary Figure 1**). Both resting state blocks were 5 minute-long recordings of EEG and electrocardiography (ECG) with eyes open. After R1, participants conducted a binary choice decision-making task with contingencies that changed over the course of learning as in previous work (Behrens et al., 2007; de Berker et al., 2016; Iglesias et al., 2013). In our task, participants completed two blocks of 200 trials each (TB1, TB2), and their goal was to find out which one of two visual icons (always either blue or orange: see **Figure 1**) would lead to a monetary reward (positive reinforcement, 5 pence). Thus, they had to learn the probability of reward assigned to each stimulus (reciprocal: p, 1-p). Both experimental blocks were divided into 5 segments with different stimulus-outcome contingency mappings that were randomly ordered for each participant and varied in length between 26 and 38 trials. These contingencies ranked from being strongly biased (90/10), moderately biased (70/30), to unbiased (50/50), and repeated in reverse relationships (10/90; 30/70) so that over the two blocks there were 10 contingency blocks in total (de Berker et al., 2016).

**Figure 1.**
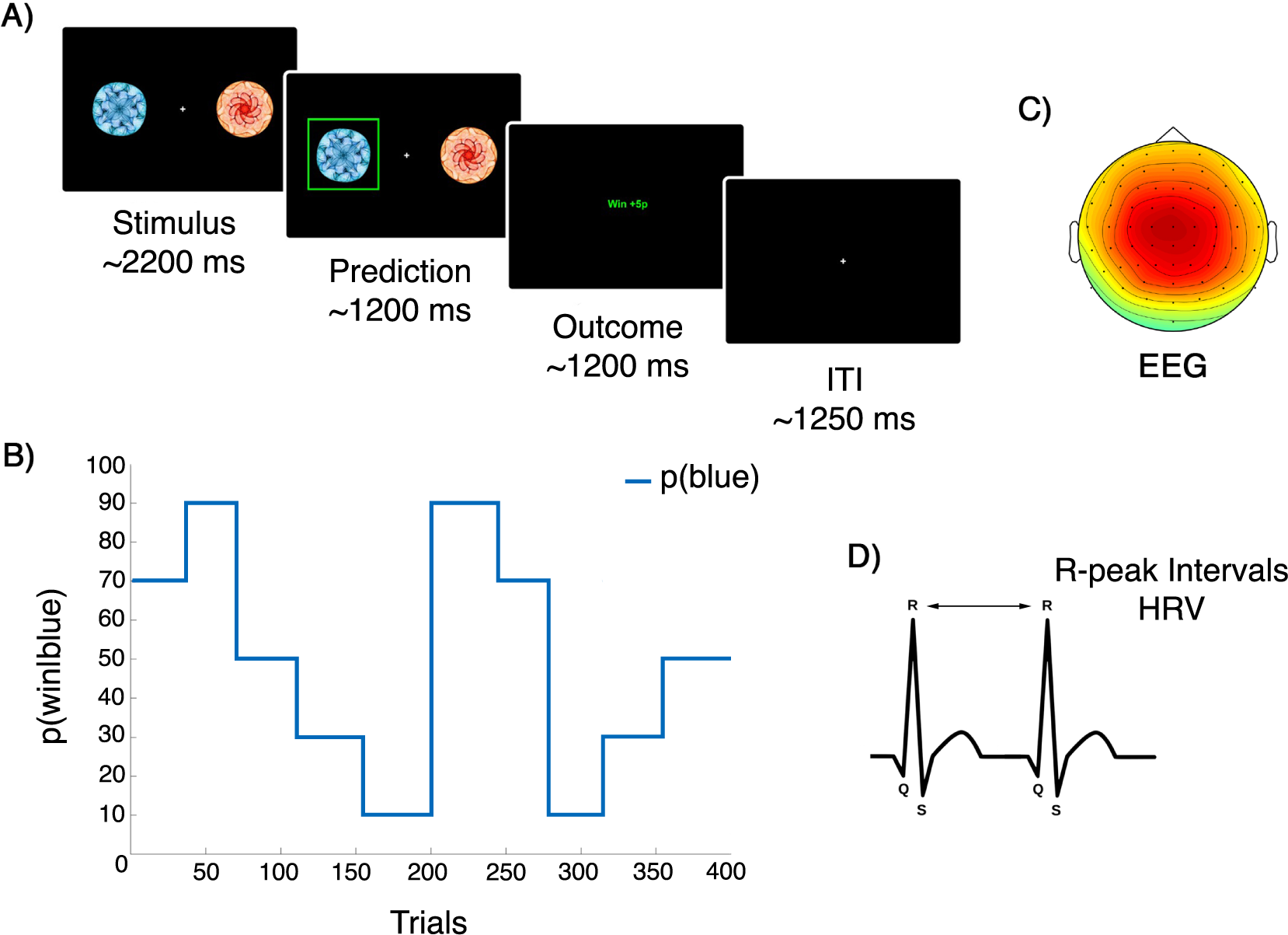
Behavioural task structure and physiological measures. A) On individual trials participants were presented with two visual icons. They were instructed to predict the rewarding stimulus (win = 5p). The stimuli (blue or orange image) were randomly presented to either the left or right of the screen. They remained on the screen until a response was given or the allowed time (2200 ms ± 200 ms) expired – recorded as no-response. When a response of either the left arrow key or right arrow key was pressed, participants immediately saw their chosen image highlighted in bright green, which remained on screen for 1200 ms (±200 ms) before the outcome was revealed. The outcome, either win or lose, was shown in the middle of the screen for 1200 ms (±200 ms) in green and red respectively. Each trial ended with a fixation cross and an inter-trial interval of 1250 ms (±250 ms). **B)** The probability governing the likelihood of the blue stimulus being rewarded (p(win|blue), with reciprocal probability values for the orange stimulus: p(win|orange) = 1 - p(win|blue)). Probability mappings varied in length (26-38 trials) ranging from heavily biased (90/10), moderately biased (70/30), to unbiased (50/50), and repeated in reverse relationships (10/90; 30/70). Here we follow one example of contingency changes for p(win|blue) over the course of the experimental blocks (TB1, TB2, 200 trials each). These blocks were divided into the 5 randomly ordered stimulus-outcome mappings and were randomly generated for each participant. While conducting the experimental task, participants’ physiological responses - **C)** EEG and **D)** ECG - were recorded continuously, with R-peaks from ECG signals being used to calculate heart-rate variability.

On individual trials, participants were asked to predict which of the two visual icons was going to reward them with money. Successful predictions were rewarded 5p, while unsuccessful predictions and no-responses were regarded as losses with 0p reward (**Figure 1**). The stimuli were either presented to the left or right of centre screen randomly. They remained on the screen until a response was given or the prediction time (2200 ms ±200 ms) expired. When a response of either the left arrow key or right arrow key was pressed, participants immediately saw their chosen image highlighted in bright green, which remained on screen for 1200 ms (±200 ms) before the outcome was revealed. The outcome, either win or lose, was shown in the middle of the screen for 1200 ms (±200 ms) in green and red respectively. Each trial ended with a fixation cross and an inter-trial interval of 1250 ms (± 250 ms).

The participants were given full computerised instructions for each element of the experiment, including questionnaires. Each questionnaire came with written instructions and was responded to using the numerical keyboard buttons. Just before 10 practice trials of the same probabilistic reward-learning task used in the main experiment, participants were explicitly informed that the reward structure would change throughout the task and that they needed to adjust their predictions in response to inferred changes (de Berker et al., 2016). Importantly, directly after this but before TB1, all were informed that this experiment was, in fact, an examination of performance using two tasks, reward-learning and public speaking; participants were instructed according to their group allocation in StA or Cont.

### 2.3 State Anxiety Manipulation

Those participants in StA were informed that they had been randomly selected to complete a public speaking task after finalising the reward-learning task (Feldman et al., 2004; Lang et al., 2015). Participants were told they would be required to present a piece of abstract art and would be allowed to prepare for 3 minutes for a 5 minute presentation of this artwork to a panel of academic experts. Those in the control group (Cont), were informed that they were to be given a piece of abstract art and they were to describe it to themselves (instead of a panel of experts) for the same period of time. After completion of the reward-based learning blocks, participants in the StA group were informed about the sudden unavailability of the panel, and thus were instructed to present the artwork to themselves (similarly to the Cont group).

### 2.4 EEG and ECG Recording and Pre-Processing

EEG and ECG signals were recorded throughout all task blocks (R1, TB1, TB2, and R2) using the BioSemi ActiveTwo system (64 electrodes, extended international 10–20) with a sampling rate of 512 Hz. The EEG signals were referenced to two electrodes affixed to the left and right earlobes. Four additional external electrodes in a bipolar configuration were used, which included two electrodes positioned to capture vertical and horizontal eye-movements (EOG), one to the zygomatic bone of the right eye, and one to the glabella (between both eyes); and two electrodes to record the ECG. ECG electrodes were placed in a two-lead configuration (Moody and Mark, 1982) calibrated to fit the Einthoven triangle (Wilson et al., 1931). All electrodes used highly conductive bacteriostatic Signa gel (by Parker). All events, including presentation of stimuli, participant responses, and trial outcomes were recorded in the EEG file using event markers.

Analysis of the ECG data was conducted in MATLAB (The MathWorks, Inc., MA, USA) using the FieldTrip toolbox (Oostenveld et al., 2011) and their recommended procedure to detect the cardiac artefacts (http://www.fieldtriptoolbox.org/example/use_independent_component_analysis_ica_to_remove_ecg_artifacts). Following this approach, the ECG signal was used to detect the QRS-complex and its main peak, the R wave peak. Next, we extracted the latency of the R-peak, which was used to compute the coefficient of variation (CV = standard deviation / mean) of the difference intervals between consecutive R-peaks (inter-beat interval). The CV of inter-beat intervals was calculated within each task block (R1: baseline, TB1, TB2, R2), and was used as a metric of heart rate variability (HRV) for statistical testing.

EEG data were preprocessed in EEGLAB toolbox (Delorme and Makeig, 2004) by first high-pass filtering at 0.5Hz (hamming windowed sinc finite impulse response [FIR] filter, 3381 points) and then notch-filtering between 48-52Hz (847 points) to remove power line noise. Afterwards, artefacts (eye blinks, eye movements, cardiac artefacts) were classified using independent components analysis (ICA, runICA algorithm) and removed (on average 2.3, SEM 0.16, components). Noisy channels were corrected utilising spherical interpolation. All signals were then epoched around outcome onsets (win, lose) from −100 to 500 ms. Noisy epochs exceeding +/−100*μV* were identified and removed using a thresholding technique relative to the pre-stimulus baseline. The number of rejected trials for each participant did not exceed 10% of the total number.

Cleaned EEG and preprocessed behavioural data files are available in the Open Science Framework Data Repository: https://osf.io/b4qkp/. The results shown in Figures 3, 4, and 5 are based on these data.

**Figure 3.**
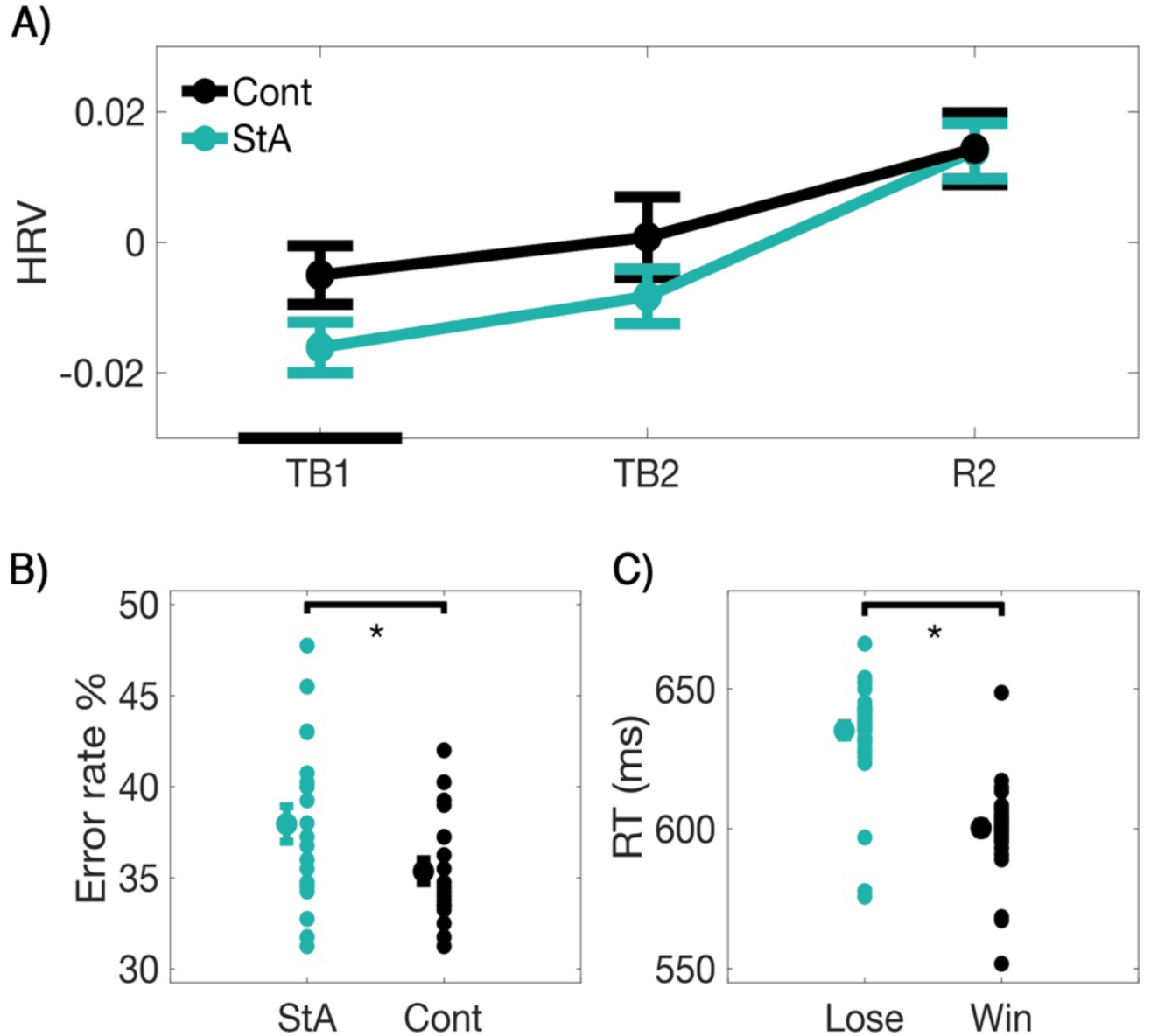
State anxiety modulates heart rate variability and behavioural responses. A) Modification in heart-rate variability (HRV) by the anxiety manipulation. The average HRV (measured with the coefficient of variation of the inter-beat-interval of the ECG signal) is provided for the state anxiety (StA) and Control (Cont) groups across task block 1 (TB1), task block 2 (TB2) and final resting state (R2). The average of the resting state (R1: baseline) has been subtracted from each subsequent task block to normalise HRV values. Significant between-group differences are identified by black bars on the x-axis (paired permutation test, *P_FDR_* < 0.05 after control of the FDR at level q = 0.05). **B)** The effect of anxiety on reward-based learning performance: error rates. Here, the average error rates of each group, the state anxiety (StA) and the control group (Cont), are presented using a central point flanked by SEM bars. To the right of each mean and SEM are the individual data points in each group to show group population dispersion. Anxiety significantly increased the error rate in the StA group when compared to Controls (*P* = 0.001). **C)** The main effect of outcome (win, black; lose, green) on mean reaction times (RT: *P* = 0). On the left the average RT of each outcome is presented using a central point with SEM bars. To the right of each mean and SEM are the individual data points of each group to show group population dispersion.

### 2.5 Measures of State Anxiety

Two markers of state anxiety were used during the experiment. First, we used the CV of the inter-beat intervals to assess HRV, as this measure, similarly to other metrics of HRV, has been reported to show reductions during anxious states (Chalmers et al., 2014; Friedman and Thayer, 1998; Gorman and Sloan, 2000; Kawachi et al., 1995). A lower HRV is associated with complexity reduction in physiological responses to stress and anxiety (Friedman, 2007; Gorman and Sloan, 2000), and is used as a transdiagnostic marker to identify anxiety in psychiatry (Quintana et al., 2016). In addition, we acquired subjective self-reported measures of state anxiety (STAI state scale X1, 20 items: Spielberger, 1983). This score was acquired twice, once before R1 (prior to the anxiety manipulation for the StA group), and once after completing the reward-learning task (just before the scheduled public speaking in the StA group). The latter score was expected to be higher in the StA group relative to the baseline pre-R1 score.

### 2.6 The Hierarchical Gaussian Filter (HGF)

We used the Hierarchical Gaussian Filter (HGF) from Mathys et al. (2014, 2011) to estimate each participant’s individual learning characteristics and belief trajectories during our binary reward-learning task. The HGF has been applied to understand learning across diverse settings (de Berker et al., 2016; Diaconescu et al., 2017b, 2014; Iglesias et al., 2013; Marshall et al., 2016; Stefanics et al., 2018; Weber et al., 2019). It is implemented in the freely available open source software TAPAS (http://www.translationalneuromodeling.org/tapas).

The HGF is a generative model representing an approximately Bayesian observer estimating hidden states in the environment. As such, the HGF is a model of perceptual inference and learning, which can be coupled to a response model. In the generative model, a sequence of hidden states *x_1_^(k)^, x_2_^(k)^,…, x_n_^(k)^* gives rise to sensory inputs that each participant encounters across *k* trials. Inference from observations to beliefs is implemented as a hierarchical belief updating process. Notably, while the perceptual model specifies how the sensory inputs are used to estimate the hidden states, the response model generates the most probable response according to those estimates (see **Figure 2**).

**Figure 2.**
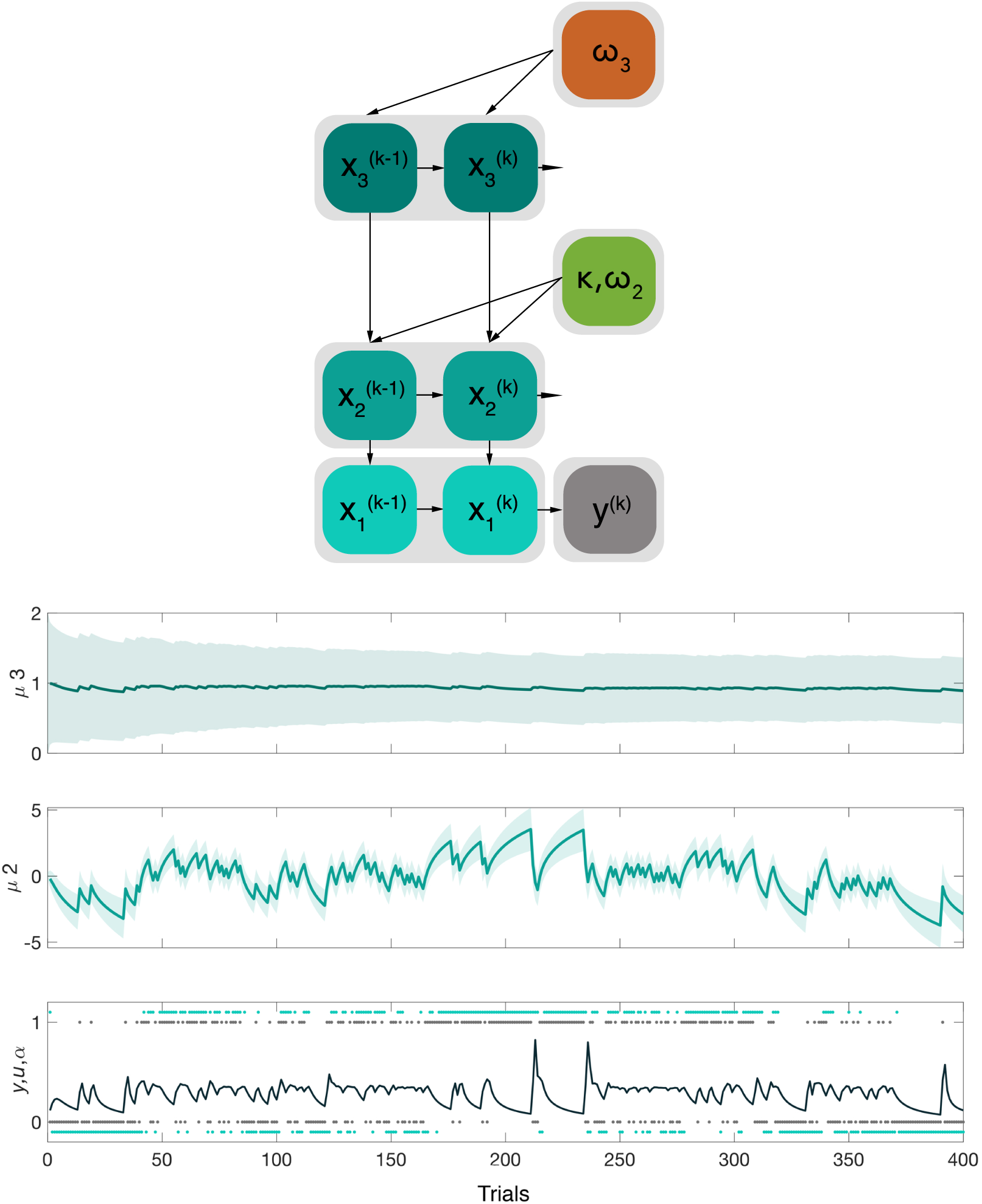
Three level binary Hierarchical Gaussian Filter for binary outcomes. Bottom panel. Representation of the three levels of the HGF for binary outcomes and the associated belief trajectories across the total 400 trials in a representative participant. At the lowest level, the inputs *u* correspond to the rewarded outcome of each trial (1 = blue, 0 = orange; shown as black dots). The participant’s responses *y* are shown in light blue dots tracking those trial outcomes. The learning rate (α) about stimulus outcomes at the lowest level is also given in black. The belief on the second level, μ_2_, represents the participant’s estimate of the stimulus tendency *x_2_* and the step size or variance of the Gaussian random walk for *x_2_* depends on parameters κ and ω_2_, in addition to the estimates of the level above*, x_3_*. The belief on the third level, μ_3_, represents estimates of volatility *x_3_*, whose step size is governed by parameter ω_3._ **Top panel.** Schematic representation of the 3-level HGF model with relevant parameters modulating each level. All parameters are fitted to individual responses of the participants and describe an individual’s learning fingerprint.

Here, we used a 3-level HGF model for binary outcomes (Mathys et al., 2014, 2011). At the lowest level, the hidden state x_1_ corresponds to the binary categorical variable of the experimental stimuli, which represents whether the blue symbol is rewarding (x_1_^(k)^ = 1; hence, orange would be non-rewarding) or not rewarding (x_1_^(k)^ = 0; with orange rewarded) in trial *k*. The second and third level states, x_2_ and x_3_, are continuous variables evolving as coupled Gaussian random walks. Thus, their value at trial *k* will be normally distributed around their previous value at trial *k-1.* State x_2_ describes the true value of the tendency of the stimulus-outcome contingency, whereas μ_2_ denotes each participant’s estimation (mean; σ_2_ being the variance) of the tendency for the probabilistic outcomes. State x_2_ can be mapped to the probability of the binary state x_1_ through a Bernoulli distribution, p(x_1_ | x_2_) = Bernoulli (x_1;_ s(x_2_)), where s(x) is a sigmoid function s(x) = 1/(1 + exp(-x)). The implied learning rate at the lowest level, α, can be defined as the change in expectation, defined as the sigmoid transformed difference between μ_2_ before seeing the input and after seeing it, relative to the difference between the observed inputs u and its prediction s(μ_2_) (**Figure 2**, lower panel; TAPAS toolbox: tapas_hgf_binary.m). A larger belief update in response to the same observed mismatch between the input u and the prediction amounts to a higher learning rate α. At the top level, x_3_ represents the phasic log-volatility within the task environment (change in the probabilistic relationships across the experiment) and μ_3_ (σ_3_) the individual’s estimate of it. The coupling between levels 2 and 3 is through a positive (exponential) function of x_3_, which represents the variance or step size of the Gaussian random walk that determines how x_2_ evolves in time:

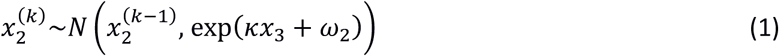

The parameters κ and ω_2_ represent the coupling strength and the tonic volatility, respectively. In the associated belief updates, momentarily high volatility estimates (μ_3_) increase the speed with which beliefs at level 2 change. Larger values of the tonic (time-invariant) part of the variance, ω_2_, generally increase the step size of x_2_ and lead to faster belief updates on level 2 irrespective of current levels of (estimated) volatility. The step size of the volatility state, x_3_, is fixed to a constant parameter ω_3_, with ω_3_ also being estimated in each individual participant, similarly to κ and ω_2_:

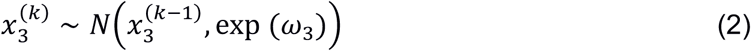

As response model we used the unit-square sigmoid observation model for binary responses (Iglesias et al., 2013; Mathys et al., 2014). This transforms the predicted probability *m(k)* that the stimulus (e.g. blue) is rewarding on trial *k* (outcome = 1) – which is a function of the current beliefs - into the probabilities p(y^(k)^ = 1) and p(y^(k)^ = 0) that the participant will choose that stimulus (blue, 1) or the alternative (orange, 0):

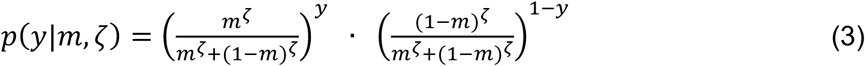

Higher values of the response parameter ζ lead to the participants being more likely to choose the response that corresponds with their current belief about the rewarded stimulus.

Fitting the combination of perceptual and response model to an individual participant’s responses allows for a subject-specific characterisation of learning (and response) style by the set of perceptual (and response) parameters. Here, we only estimated ω_2,_ ω_3,_ and ζ, with κ and the starting values of the beliefs fixed according to **Table 1**.

**Table 1.** Means and variances of the priors on perceptual parameters and starting values of the beliefs of the HGF.

| Prior | Mean | Variance |
| --- | --- | --- |
| $\kappa$ | 1 | 0 |
| $\omega_2$ | -4 | 0.5 |
| $\omega_3$ | -7 | 0.5 |
| $\mu_2^0$ | 0 | 0 |
| $\sigma_2^0$ | $\log(0.1)$ | 0 |
| $\mu_3^0$ | 1 | 0 |
| $\sigma_3^0$ | $\log(1)$ | 0 |
| $\zeta$ | $\log(48)$ | 1 |

Importantly, the update equations of the posterior estimates for level *i (i* = 2 and 3) depend on the prediction error of the level below, δ_i-1,_ scaled proportionally to the ratio of the precision of the prediction of the level below (hat denotes prediction before seeing the input) and the precision of the current level. This is captured in the expression:

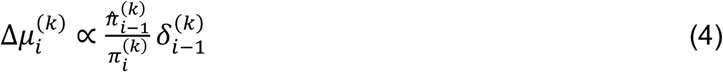

And precision is defined as the inverse variance of the expectation:

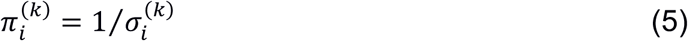

The variance of the posterior expectation, σ_i_, corresponds to the estimation or informational uncertainty about the hidden state x_i_. Accordingly, equation 4 above articulates the idea that more uncertain (less precise) belief estimates for the current level should motivate larger changes to beliefs. More detailed update equations for our 3-level HGF model are supplied in Mathys et al. (2011, 2014). The additional measure of uncertainty that we used was environmental uncertainty, which is related to volatility in the environment, according to this expression:

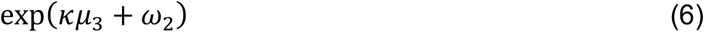

In sum, in the current study, the computational quantities of interest were the model parameters ω_2_ (tonic volatility estimate) and ω_3_ (‘meta-volatility’); the trial-wise posterior beliefs about volatility (μ_3_) – which were used to estimate trial-wise environmental uncertainty; and the trial-wise variances on levels 2 and 3 (σ_2_, σ_3_) as a measure of (informational) uncertainty about the hidden states on these levels.

Because precision-weighted prediction errors play an important role in current Bayesian theories of perceptual inference and learning (Doya et al., 2007; Feldman and Friston, 2010; Friston et al., 2013; Friston and Kiebel, 2009; Moran et al., 2013; Rao and Ballard, 1999), and these are the quantities that are considered to predominantly modulate EEG signals (Friston and Kiebel, 2009), we selected the pwPE trajectories from levels 2 and 3 (labelled ε_2_, ε_3_) to examine how these are represented in the brain as a function of state anxiety (see GLM analysis section below).

### 2.7 Model Space

We used four computational models of learning. The first two were a 2-level (excluding volatility) and 3-level hierarchical Bayesian model (HGF: Mathys et al. (2011). The third model was a Rescorla Wagner (RW) where PEs drive belief updating but with a set learning rate (Rescorla and Wagner, 1972). The final model was a Sutton K1 model (SK1) that permits the learning rate to change with recent prediction errors (Sutton, 1992). These models are also implemented in the TAPAS toolbox. Models were then compared at the group level for fit using random effects Bayesian model selection (BMS; Stephan et al., 2009; code from the freely available MACS toolbox; Soch and Allefeld, 2018). BMS provided model frequencies and exceedance probabilities reflecting how optimal each model or family of models performed (Soch et al., 2016). First, the log-model evidence (LME) from both Bayesian models were combined to get the log-family evidence (LFE) and was compared to the LFE of the family of reinforcement learning models (RW and SK1) to assess which provided more evidence. In the winner family, additional BMS determined the final optimal model.

### 2.8 EEG analysis and the General Linear Model

Prior to single-trial ERP analysis using the general linear model (GLM), a statistical analysis of the differences between ERPs following win versus loss outcomes was conducted independently in both groups (StA, N = 21, Cont, N = 21) using permutation tests with a cluster-based threshold correction to control the family-wise error (FWE) at level 0.05 (dependent samples t-test, 1000 iterations; (Maris and Oostenveld, 2007); FieldTrip toolbox, (Oostenveld et al., 2011). Experimental cluster-based test statistics being in the 2.5th and 97.5th percentiles of the permutation distribution were considered significant (two-tailed test, *P* < 0.025). For this statistical analysis, the ERP data epochs were baseline-corrected by subtracting the mean activation during the baseline period from −200 ms to 0 ms. The aim of this within-group ERP analysis was to assess whether the windows associated with the effect of the outcome (win, lose) on the EEG signals in our task and in each group separately converge with the windows of the fERN and P300 effects reported in previous studies (see for instance Nieuwenhuis et al., 2004; Hajcak et al., 2005). Note that the windows selected for the GLM analysis were broadly based on the temporal intervals of the fERN and P300 components, although pwPE regressors can actually modulate the trial-wise ERP responses peaking at different latencies than the model-free ERP effects (see e.g. Diaconescu et al. [2017a], Weber et al. [2019], Stefanics et al. [2018]).

For the GLM single-trial analysis, EEG data were downsampled to 256 Hz, low-pass filtered at 30Hz and converted to SPM 12 (http://www.fil.ion.ucl.ac.uk/spm/ version 7487) (Penny et al., 2011). In SPM 12 software we converted the EEG data into 3-dimensional volumes (two spatial dimensions: anterior to posterior, left to right across the scalp; one temporal dimension: peri-stimulus time; Litvak et al., 2011). All participants’ data comprised of 64 channels and 78 time points using a voxel size of 4.2 mm × 5.4 mm × 4 ms and were spatially smoothed to adjust for between-subject spatial variability in the channel space. In accordance with fMRI, the scalp x time 3D images were then tested statistically using statistical parametric mapping and the GLM (see next section; Kiebel and Friston, 2004a, 2004b; Kilner and Friston, 2010). This procedure is firmly established in EEG using SPM (Litvak et al., 2011).

Our GLM was composed of trial-wise estimates of two computational quantities: absolute values of pwPEs in level 2 (ε_2_), and pwPEs in level 3 (ε_3_). The absolute value of ε_2_ was selected because its sign is arbitrary: the quantity x_2_ is related to the tendency of one choice (e.g. blue stimulus) to be rewarding (x_1_ = 1), yet this choice was arbitrary and thus is the sign of the pwPE at this level (see for instance Stefanics et al., 2018). In addition, we used as third regressor the trial-wise outcome values (0 for lose, 1 for win) as we expected this variable to account for much of the signal variance in the EEG epochs. These three regressors were not orthogonalised. The window for this analysis was selected from 200 to 500 ms, based on previous literature on the fERN (also ERN) and P300 components (Hajcak et al., 2005; Nieuwenhuis et al., 2004).

Using these choices for regressors and time interval, we then carried out a whole-volume (spatiotemporal) analysis that searched for representations of our computational quantities in the single-trial EEG responses for each individual participant, before assessing within-group statistical effects at the second level. We corrected for multiple comparisons across the whole time-sensor matrix using Gaussian random field theory (Worsley et al., 1996) with a family-wise error (FWE) correction at the cluster-level (p<0.05). This was performed with a cluster defining threshold (CDT) of p<0,001 (Flandin and Friston, 2019). Importantly, all results reported survived whole-volume correction at the peak-level (p<0.05). We assessed separately within each group whether the trajectories of our computational quantities were associated with increases or decreases in EEG amplitudes using an F-test. A standard summary statistics approach was used to perform random effects group analysis within each group (StA, Cont) of 21 participants independently (Penny and Holmes, 2007). As we hypothesised between-group differences in the uncertainty estimates, which would differently affect the precision weights on the PEs, thus the ε_2_ and ε_3_ regressors, we did not implement a between-group statistical analysis on the GLM-driven EEG representations as this would result in invalid statistical inference (Kriegeskorte et al., 2009).

### 2.9 Statistics

To assess Group (StA, Cont) and Block (1,2) main effects and interactions in state anxiety measures, behavioural and computational model variables, we applied non-parametric factorial synchronised permutations tests (Basso et al., 2007). These permutation-based factorial analyses were followed up by planned pair-wise permutation tests to assess our specific hypothesis of between-group differences. This applies to the following dependent variables: (a) model-free behavioral measures (error rate, reaction time: RT); (b) CV as a measure of HRV (CV values in TB1, TB2 and R2 blocks were corrected by subtracting the R1 baseline value); (c) HGF parameters (tonic volatility ω_2_ modulating the variance of the Gaussian random walk at level 2; and ω_3_, the step size at the highest level); (d) HGF estimates of belief trajectories (informational uncertainty about x_2_ [σ_2_], environmental uncertainty, related to volatility in the environment: exp(κμ_3_ + ω_2_), and uncertainty about volatility x_3_ [σ_3_]).

Pair-wise permutation tests were also used to test within-group differences in RT across blocks. In the case of multiple comparisons (for instance, two between-group permutation tests run separately for each block), we controlled the false discovery rate (FDR) using an adaptive linear step-up procedure set to a level of q = 0.05 (Benjamini et al., 2006). This procedure furnished us with an adapted threshold p-value (P_FDR_). Prior to these statistical analyses and following BMS, the trial-wise trajectory for each computational quantity of interest (σ_2_, σ_3_ or environmental uncertainty, eq. 6) was extracted from the winning model, followed by an average across trials within task blocks (TB1, TB2). Despite of trial-by-trial changes in these belief trajectories relating to the subject-specific trial structure and contingency blocks, the average values across trials revealed the general monotonic changes in the trajectories within each block, which is what we aimed to evaluate as a function of the factors Group and Block using the 2 x 2 factorial analysis, as described above.

Below in the Results section, we present the mean and standard error of the mean (SEM) for our dependent variables (either in text or in a figure), alongside non-parametric effect sizes for pair-wise comparisons and corresponding bootstrapped confidence intervals (Grissom and Kim, 2012; Ruscio and Mullen, 2012). In the case of within-group comparisons, the non-parametric effect size was estimated using the probability of superiority for dependent samples (Δ*_dep_*), whereas for between-group effects we used the probability of superiority (Δ); both are calculated in line with Grissom and Kim (2012), expressed as the number of values in sample A greater than those in sample B (Δ = P[A>B]). In the case of dependent samples, the comparison between pairs is done for matched pairs. Although in the original formulation by Grissom and Kim (2012), ties were not taken into account; here, in line with Ruscio and Mullen (2012), we corrected (Δ) using the number of ties (difference scores = 0) and estimated bootstrapped confidence intervals (CI) for (Δ).

## 3. Results

### 3.1 Heart-rate variability

Using a non-parametric 2 x 3 factorial test with synchronised rearrangements, significant main effects of Block (*P* = 0.0016) and Group (*P* = 0.0042) were revealed. No interaction effect was found. Planned between-group comparisons using permutation testing revealed significantly lower HRV in StA during TB1 (mean 0.02, SEM 0.003) when compared to Cont (mean 0.01, SEM 0.004, *P_FDR_* < 0.05, Δ = 0.70, CI = [0.54, 0.86], see **Figure 3A**). These results indicate that the experimental manipulation achieved physiological changes from the heart corresponding to an anxious state (Chalmers et al., 2014; Feldman et al., 2004).

### 3.2 State-trait Inventory

Self-reported state anxiety measures led to a significant main effect of Block (*P* = 0.03). There were no significant main effect of Group or interaction effect (*P* = 0.54, *P* = 0.85). Assessing further the main Block effect in each group separately, we observed a significant increase in self-reported state anxiety in StA from just before TB1 (mean 36.3, SEM 1.99) to after TB2 (mean 40.6, SEM 2.42, *P* = 0.002, Δ_dep_ = 0.64, CI = [0.56, 0.71]). This was also significant for Cont, with self-reported state anxiety just before TB1 (mean 35.0, SEM 1.54) increasing after TB2 (mean 38.9, SEM 1.54, *P* = 0.001, Δ_dep_ = 0.67, CI = [0.58, 0.77]). No difference in trait anxiety scores between groups was found (*P* > 0.1).

### 3.3 Model-free Analysis

The percentage of errors made by each participant across 400 trials was used as a measure to assess whether anxiety impairs reward-learning task performance. Using non-parametric factorial test (synchronised rearrangements), the main effect of factor Group on error rates was significant (*P* = 0.01), but not the main effect of task Block or interaction effect (*P* = 0.056, *P* = 0.662). Planned between-group comparisons using pair-wise permutation tests revealed a significantly higher total average error rate in the StA group (mean 38.0, SEM 0.97), in comparison to the Cont group (mean 35.6, SEM 0.66, *P* = 0.001, Δ = 0.70, CI = [0.58, 0.82], see **Figure 3B**).

Turning to the mean reaction times (RT, in milliseconds), a significant main effect of task Block was observed (*P* = 0.008). But no significant main effect of Group or interaction effect was found (*P* = 0.64, *P* = 0.26) in line with previous work on anxiety (Bishop, 2009). Post-hoc analyses on the Block effect in each group revealed there was a significant decrease in the mean RT from TB1 (mean 658.3, SEM 32.53) to TB2 (mean 552.8, SEM 20.68), in the StA group: *P* = 0, Δ = 0.73, CI = [0.65, 0.81]. This effect was also significant for Cont, with mean RT dropping from TB1 (mean 657.7, SEM 35.65) to TB2 (mean 591.4, SEM 31.78, *P* = 0.005, Δ = 0.65, CI = [0.56, 0.72]). As a final analysis, we contrasted the mean RT across all participants between outcomes on lose trials (mean 639.8 ms, SEM 0.36) and win trials (mean 600.8 ms, SEM 0.33). This was highly significant, as expected, reflecting a slower response in lose trials (*P* = 0, Δ = 1, CI = [1, 1], **Figure 3C**).

### 3.4 Bayesian Model Selection

After fitting each model (HGF: 3-Levels, 2-Levels, the Rescorla Wagner [RW], and Sutton K1 [SK1]) individually in each of the 42 participants and obtaining log-model evidence (LME) values for each, we compared the four models using Bayesian model selection (BMS). Results from BMS revealed that the family of Bayesian models (3-levels and 2-levels HGF) had much stronger evidence than the reinforcement-learning models (RW, SK1), with an exceedance probability of 0.99, and an expected frequency of 0.73 (leftmost columns: **Figure 4A**). Next, within the Bayesian models, an additional BMS step using the LME for each subject and model demonstrated much stronger evidence for the 3-levels HGF model relative to the 2-levels version, with an exceedance probability of 0.98 and an expected frequency of 0.68 (rightmost columns: **Figure 4A**). The 3-levels HGF model was the winner model also when performing BMS separately in the StA and Control groups.

**Figure 4.**
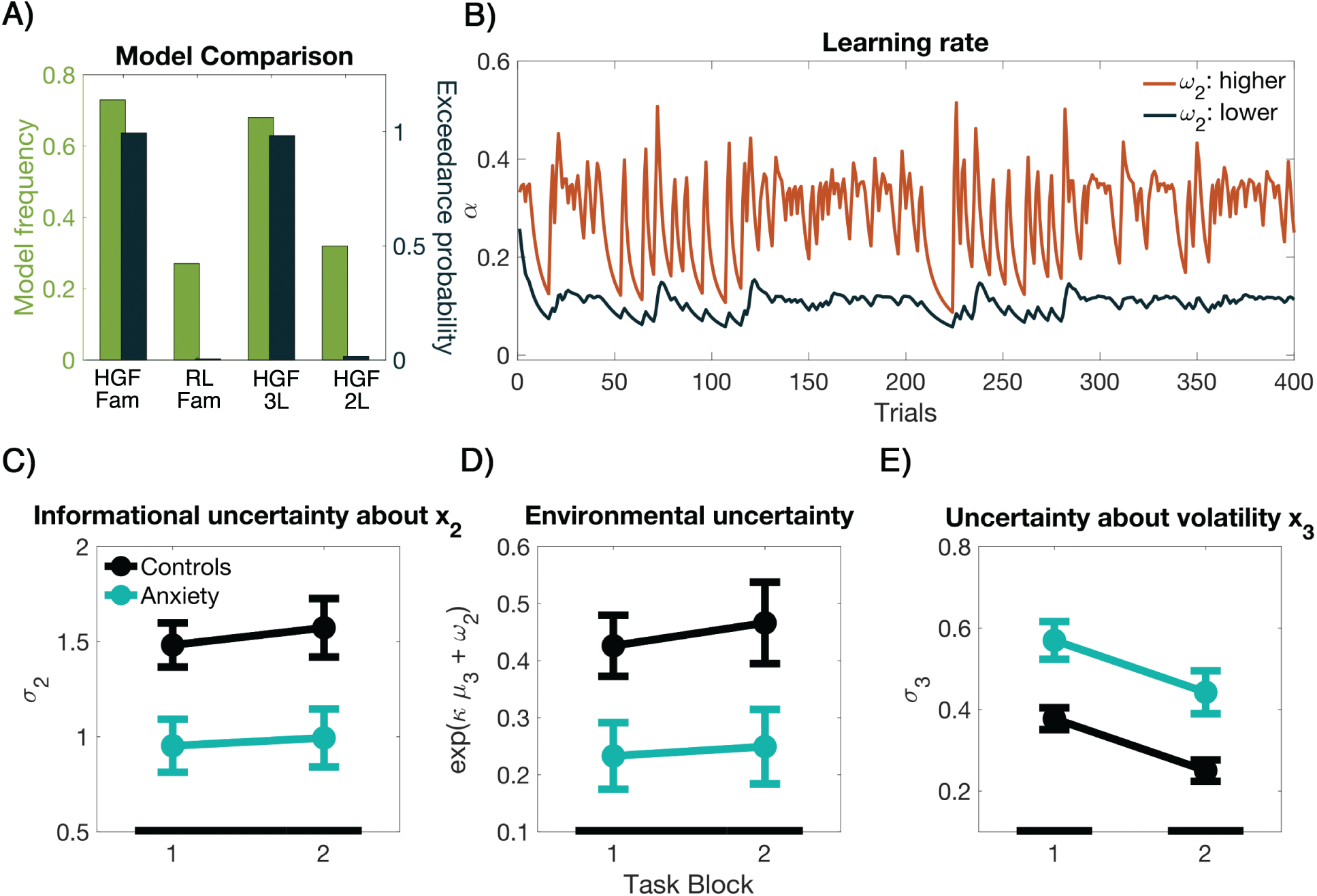
Bayesian Model Selection and Hierarchical Gaussian Filter Results. A) Bayesian model selection (BMS). The two leftmost columns represent the model frequency (green) and exceedance probability (black) for the family of models ‘HGF Fam’ (3-Level and 2-Level HGF) and the family of reinforcement learning models ‘RL FAM’ (RW, SK1). The family of HGF models provided the best model evidence. In the two right columns the comparison between the two HGF models is shown (3-Level: HGF 3L and the 2-Level: HGF 2L). The 3-Level HGF provided stronger model evidence. **B)** Illustration of the trial-by-trial learning rate about stimulus outcomes (α) in two ideal observers with different values of ω_2_. Trajectories were simulated using the same input sequence and parameters (except ω_2_): μ_2(0)_ = 0, μ_3(0)_ = 1, σ_2(0)_ = log(0.1), σ_3(0)_ = log(1), κ = 1, ω_3_ = 7. The two priors on ω_2_ used in the simulated trajectories are −2 (orange) and −4 (black). This parameter represents the tonic part of the variance in the Gaussian random walk for x_2_ and modulates the learning rate about stimulus outcomes at the lowest level. Lower ω_2_ values lead to smaller trial-by-trial learning increments. When comparing ω_2_ values between groups (StA, Cont), we found more negative values in StA than in the Cont group (*P* = 0.002). **C)** Lower ω_2_ in state anxiety leads to decreased informational uncertainty about x_2_. There was a significant main effect for factor Group (StA, green; Cont, black; synchronised permutation test: P = 0.003) but not for factor Block (*P* > 0.05). Planned between-group comparisons indicated that state anxiety significantly decreased the average uncertainty about beliefs on tendency x_2_ (*P* = 0.003, as given by black bars), after averaging across both blocks; significant effect indicated by black bars at the bottom). **D)** Lower ω_2_ in state anxiety leads to decreased environmental uncertainty (*P* = 0.02) (not effect of factor Block (*P* > 0.05)). Thus, StA participants had a lower estimate of environmental uncertainty or volatility. **E)** State anxiety increased uncertainty about volatility in the task environment (σ_3_). We found a significant main effect for factor Block (*P* = 0.004) and Group (StA, green, Cont, black; *P* = 0.0002), modulating uncertainty about volatility. Planned between-group comparisons further indicated that state anxiety exhibited significantly higher σ_3_, as compared to control participants, separately in each task block (TB1, TB2, P_FDR_ < 0.05, as given by black bars).

### 3.5 Model-based analysis

#### 3.5.1 State anxiety is associated with a lower learning rate about stimulus outcomes

We observed significant differences between the groups in parameter ω_2_, which is the tonic part of the variance of the Gaussian random walk for x_2_, or the tonic volatility estimate. Smaller values were obtained in StA (mean −3.1, SEM 0.23) than in the Cont group (mean −2.0, SEM 0.15, *P* = 0.002, effect size: Δ = 0.75, CI = [0.55, 0.90]). No differences were found in ω_3_, the log-variance of at the top level. The values of ω_2_ influence, among other HGF trajectories, the learning rate at the lowest level, α (through a modulation of μ_2_), driving the step of the update about stimulus outcomes (Mathys et al., 2014). More negative ω_2_ values – as found in StA – lead to smaller updates, and thus to smaller learning rates (See illustration in **Figure 4B).**

#### 3.5.2 Informational Uncertainty about the outcome tendency is lower in state anxious individuals

We then evaluated the model estimates of informational uncertainty about the outcome tendency, σ_2_. This variance measure reflects the lack of knowledge about x_2_ and depends on ω_2_ but also on the volatility estimate μ_3_ and other quantities (eqs. 11 and 13 in Mathys et al., 2014). Thus, lower ω_2_ values lead to smaller σ_2_, however, the impact of μ_3_ on σ_2_ could alter this effect. Here we found a significant main effect of Group (*P* = 0.003). Yet, the Block factor and interaction effect were not significant (*P* = 0.4028, *P* = 0.7352). In addition, planned comparisons showed that anxiety significantly lowered the total average σ_2_ for StA in comparison to Cont (**Figure 4C**; *P* = 0.003, Δ = 0.75, CI = [0.55, 0.89]). Because precision is the inverse variance (informational uncertainty) of the distribution, these outcomes demonstrate that StA individuals estimated their beliefs about the outcome tendency to be more precise, and therefore new information had a smaller impact on the update equations for x_2_.

#### 3.5.3 Environmental uncertainty is underestimated in state anxiety

Environmental uncertainty, which relates to the posterior beliefs over volatility, depends only on the tonic volatility, ω_2_, and the trial-wise volatility estimate, μ3 (see equation 6 above; the coupling constant κ was fixed to zero). We found that environmental uncertainty was significantly modulated by factor Group (*P* = 0.02), while there was no significant main effect for factor Block or interaction effect (*P* = 0.58, *P* = 0.7547). Further pair-wise analyses demonstrated that the StA group underestimated the environmental uncertainty, relative to control participants, when averaging across both experimental blocks (**Figure 4D**; *P = 0*, Δ = 0.74, CI = [0.54, 0.88]).

#### 3.5.4 Uncertainty about volatility is higher in state anxious individuals

In contrast to the effect on σ_2_ reported above, state anxiety increased uncertainty on level 3 (σ_3_). We found both a significant main effect of Block (*P* = 0.0004) and Group on this parameter (*P* = 0.0002), yet no interaction effect (*P* = 0.99). Across blocks, the informational uncertainty generally decreased. Planned comparisons demonstrated that separately in the first and second task blocks anxiety significantly increased σ_3_ in the StA group when compared to the Cont group (**Figure 4E**; P_FDR_ < 0.05, effect size for TB1: Δ = 0.74, CI = [0.53, 0.88]; TB2: Δ = 0.74, CI = [0.53, 0.89]). The larger anxiety-induced informational uncertainty about volatility corresponds to less precise estimates about volatility, and thus, to a larger impact of new information on the update of volatility estimates, μ_3_.

### 3.6 Standard Lose versus Win ERP results

Cluster-based random permutation tests demonstrated in both groups (StA, 21, Cont, 21) a significant difference between the effect of the two outcomes (lose, win) on the ERP (two significant clusters in each group at level *P <* 0.025).

In the control group, losing led to a more negative ERP amplitude than winning during a time window between 230 and 360 ms post outcome (negative cluster, *P* = 0.008). This effect at first had a centro-parietal distribution, which later propagated to broader central, frontal, temporal, and parietal electrode regions, occurring approximately in line with the fERN ERP (**Supplementary Figure 2**). In a later time window, between 350 and 500 ms, losing evoked a more positive amplitude when compared to winning (positive cluster, *P* = 0.0002). During this later latency, the difference originated over fronto-central electrodes, and later spread to centro-parietal electrodes resembling the P300 component wave (**Supplementary Figure 2**). The latency of the significant clusters confirmed that lose relative to win trials elicited a biphasic ERP modulation consisting of an earlier negative wave resembling the fERN and a later positive and very pronounced deflection corresponding to the P300.

In the state anxiety group, a similar spatio-temporal ERP profile to the Cont group emerged. Losing was associated to a more negative ERP amplitude when compared to winning between 240 and 350 ms post outcome (significant negative cluster, *P* = 0.004; **Supplementary Figure 3**). This effect originated in centro-parietal regions and spread to frontal and central sites later in the time window. Following this effect, we found a significant positive deflection between 350 and 500 ms (positive cluster, *P* = 0.0002; **Supplementary Figure 3**). In this later time window, the spatio-temporal pattern began with activation across fronto-central electrodes and developed in to centro-parietal electrodes regions, like in the control group, but in more anterior central electrodes.

### 3.7 Single-trial ERP modulations by precision-weighted PEs

The HGF results had confirmed that state anxiety alters informational uncertainty estimates about beliefs on level 2 and also about volatility on level 3 (**Figure 4**), in an opposing pattern of changes (decrease in σ_2_ and increase in σ_3_ relative to control participants). We then proceeded to analyse in each group separately the electrophysiological representations of trial-wise pwPEs for level 2 and 3 – which are a function of those uncertainty estimates as shown in equation 4 (for an illustration of ε_2_, ε_3_, see **Figure 5A**). The GLM results of the additional outcome regressor are shown in **Supplementary Figure 4**.

**Figure 5.**
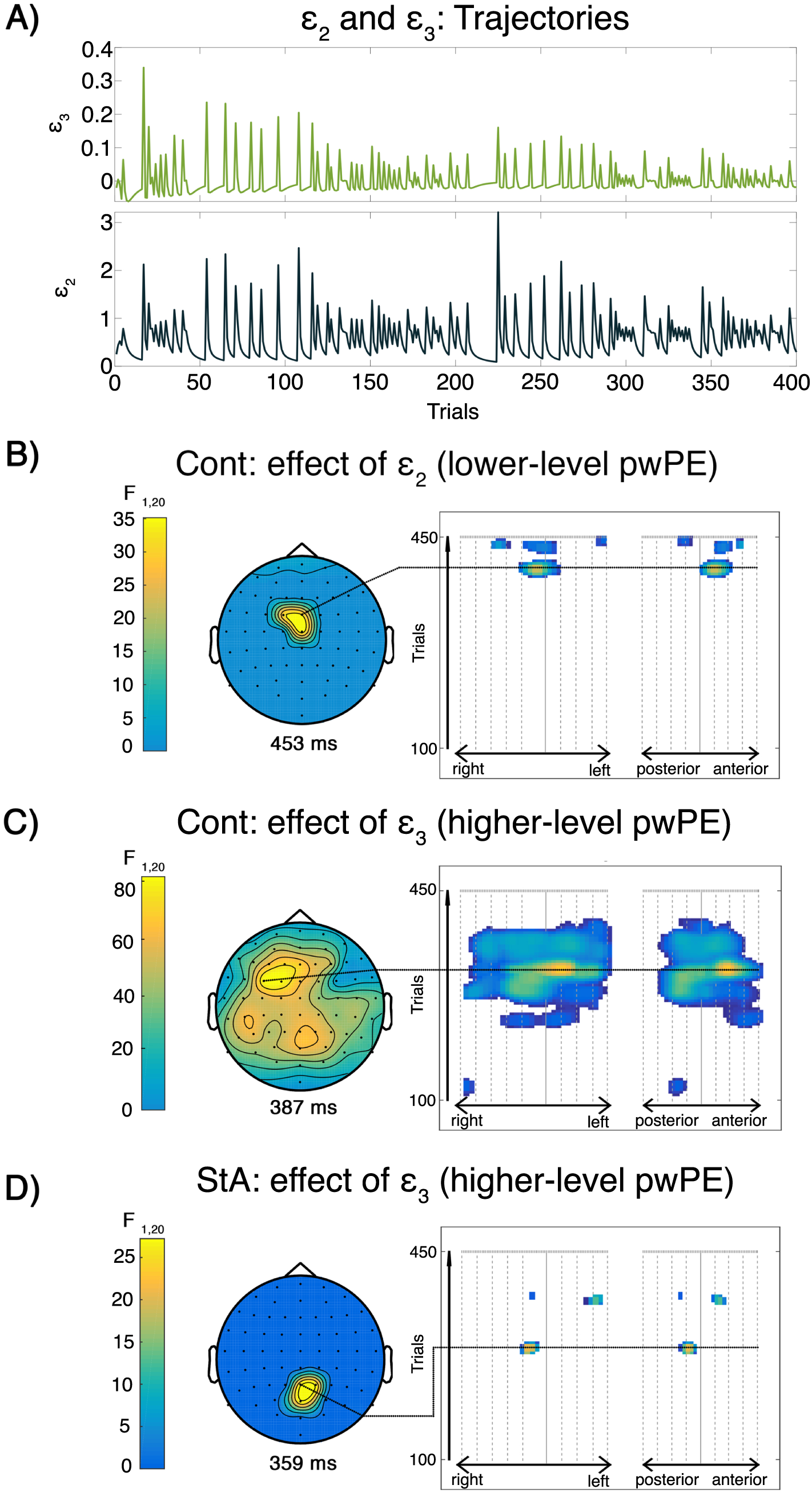
Signatures of precision-weighted prediction errors on trial-wise ERPs. A) Trajectories of model-based estimates for both lower-level and higher-level pwPE for one representative control group participant across 400 trials. In green are higher-level pwPEs concerning volatility; in black are the absolute values of the lower-level pwPE concerning beliefs about the rewarding stimulus. **B)** Effect of pwPEs on level 2 (ε2) in controls. In the control group (Cont), responses in single-trial ERPs were significantly modulated by pwPEs about reward tendency in central electrodes. This significant cluster occurred between 440 ms and 461 ms, and is shown on a 2D scalp map at the time of the maximum peak of the cluster (453 ms post stimulus, PFWE = 0.007, with a cluster-defining threshold of P < 0.001). **C)** Effect of pwPEs on level 3 (ε3) in controls. In the Cont group, pwPEs about volatility estimates correlated with in activation changes across a left frontocentral region between 304 ms to 450 ms, as shown in this topographical representation at the time of the maximum peak of the cluster (387 ms post stimulus, PFWE < 0.0001, with a cluster-defining threshold of P < 0.001). **D)** Effect of pwPEs on level 3 (ε3) during state anxiety. In the state anxiety group (StA), ε3 was associated with trial-wise ERP changes in midline parietal electrodes. This effect, ranging from 354 ms and 365 ms, is shown in a topographic scalp map at the time of the maximum peak of the cluster (359 ms post stimulus, PFWE = 0.028, with a cluster-defining threshold of P < 0.001). A further significant effect of a smaller cluster size occurred between 423-431 ms (PFWE = 0.035) at the left temporal region.

#### 3.7.1 Low-level precision-weighted prediction errors

A significant effect was found in the Cont group between 440 ms and 461 ms post stimulus, peaking at 453 ms over central channels (**Figure 5B**; whole-volume cluster-level FWE corrected, termed *P*_FWE_ hereafter, *P_FWE_* = 0.007). Two additional clusters were found slightly later around 480-490 ms at right frontal (*P_FWE_* = 0.039) and central (*P_FWE_* = 0.020) electrodes. Details on the cluster effects can be found in **Table 2**. By contrast, testing the GLM in the StA group, we found no significant modulation by ε_2_ of single-trial ERPs.

**Table 2.** Test statistics for higher and lower-level precision-weighted prediction errors and trial outcomes. Each significant activation is ordered according to size (leftmost column). We provide both the cluster and peak *p* values with the family-wise error correction applied. Also given are the relevant statistics (*F* and peak equivalent *Z*) for each activation cluster and within each activation.

| Activation size<br>(voxels) | Cluster *p* value<br>(FWE<br>corrected) | Peak *p* value<br>(FWE<br>corrected) | Peak *F*<br>statistic | Peak<br>equivalent *Z*<br>statistic | Peak<br>latency<br>(ms) |
| --- | --- | --- | --- | --- | --- |
| **Control Group**<br>- Lower-level<br>pwPE ($\epsilon_2$) | | | | | |
| 205 | 0.007 | 0.011 | 35.16 | 4.30 | 453 |
| 10 | 0.039 | 0.025 | 30.17 | 4.08 | 488 |
| 69 | 0.020 | 0.030 | 29.01 | 4.02 | 484 |
| 17 | 0.035 | 0.036 | 27.77 | 3.96 | 496 |
| **Control Group -**<br>Higher-level<br>pwPE ($\epsilon_3$) | | | | | |
| 12675 | 0.000 | 0.000 | 85.24 | 5.58 | 387 |
|  |  | 0.000 | 67.19 | 5.24 | 375 |
|  |  | 0.000 | 65.61 | 5.21 | 383 |
| 35 | 0.022 | 0.013 | 36.91 | 4.37 | 219 |
| **State Anxiety**<br>**Group**<br>- Higher-level<br>pwPE ($\epsilon_3$) | | | | | |
| 33 | 0.028 | 0.042 | 27.22 | 3.93 | 359 |
| 15 | 0.035 | 0.044 | 26.89 | 3.92 | 426 |
| 2 | 0.046 | 0.049 | 26.35 | 3.89 | 434 |
| **Control Group -**<br>Outcome |  |  |  |  |  |
| 2599 | 0.000 | 0.000 | 89.09 | 5.64 | 242 |
|  |  |  | 83.15 | 5.55 | 242 |
|  |  |  | 82.91 | 5.54 | 238 |
| 296 | 0.004 | 0.016 | 33.35 | 4.23 | 414 |
| **State Anxiety**<br>**Group**<br>- Outcome |  |  |  |  |  |
| 5260 | 0.000 | 0.001 | 50.10 | 4.82 | 293 |
|  |  | 0.001 | 49.40 | 4.80 | 277 |
|  |  | 0.004 | 41.51 | 4.54 | 246 |
| 559 | 0.003 | 0.010 | 34.70 | 4.28 | 402 |
|  |  | 0.010 | 34.54 | 4.28 | 402 |

In an attempt to understand the lack of significant effects of ε_2_ in the GLM analysis in the anxiety group, we evaluated the variance of this regressor in every participant and then calculated the mean in each group separately. In StA, the group mean variance of ε_2_ was smaller (0.14, SEM 0.048) than in the control group (0.31, SEM 0.080), suggesting that less variance in this regressor is available in the StA group to explain the EEG variance.

#### 3.7.2 High-level precision-weighted prediction errors

In the Cont group, ε_3_ significantly elicited trial-wise EEG responses from 304 ms to 450 ms post stimulus, with a maximum effect at 387 ms across a left frontocentral region (P*_FWE_* < 0.0001). Additional significant effects of a smaller cluster size were found earlier, between 205-225 ms (*P_FWE_* = 0.022) at right parietal channels (**Figure 5C**).

Precision-weighted PEs about volatility modulated significantly the ERP responses in the StA group between 354 ms and 365 ms, with a maximum at 359 ms most pronounced over midline parietal electrode regions (*P_FWE_* = 0.028, **Figure 5D**). A further significant effect of a smaller cluster size occurred between 423-431 ms (*P_FWE_* = 0.035) at the left temporal region.

## 4. Discussion

We combined computational modelling of behaviour and analysis of electrophysiological responses to examine how state anxiety modulates reward-based learning when learning in a volatile environment. Our key finding is that state anxiety was associated with a lower learning rate, driven by an underestimation of environmental and informational uncertainty. At the same time, we observed a decrease in the precision of estimates of environmental volatility – a higher-level belief – during anxiety.

Trial-wise estimates of uncertainty – or its inverse, precision – serve to scale the impact of prediction errors (PEs) on the belief updates. Consistent with previous reports (Stefanics et al., 2018; Weber et al., 2019), we found that precision-weighted PEs (pwPEs) on two hierarchical levels can explain trial-wise modulation of observed ERP responses in control participants. Specifically, lower-level pwPEs about reward outcomes explained variation in EEG amplitudes in a different time window after stimulus presentation than did higher-level pwPEs informing volatility estimates. A different pattern emerged in the state anxiety group, where only higher-level pwPEs modulated the trial-wise ERP changes. Taken together, the data suggest that temporary anxious states in healthy individuals impair reward-based learning in volatile environments, primarily through changes in uncertainty estimates and potentially a degradation of the neuronal representation of hierarchically-related pwPEs, which are considered to play a central role in current Bayesian accounts of perceptual inference and learning.

### States of anxiety bias computations of different types of uncertainty during reward-based learning

The threat of a public speaking task used in our experiment reduced heart rate variability, which is consistent with previous findings on state anxiety (Chalmers et al., 2014; Feldman et al., 2004; Gorman and Sloan, 2000), and despite the lack of corresponding significant effects in the STAI state anxiety scale. At the same time, our experimental manipulation had an adverse effect on reward-based learning. Having matched trait anxiety levels across the state anxious and the control group, our results indicate that the changes observed in reward-based learning – lower learning rates and changes in uncertainty – can be linked to temporary anxious states independent of trait levels. These outcomes thus expand prior findings of an association between high levels of trait anxiety and difficulties in decision-making tasks (de Visser et al., 2010; Miu et al., 2008) and learning in volatile task environments (Browning et al., 2015; Huang et al., 2017) to the realm of state anxiety. Moreover, using the threat of public speech, our approach allowed us to investigate behavioural, physiological, and neural responses in anticipation of a future unpredictable threat. Aberrant anticipatory responding to upcoming uncertain threats has been proposed to be a common explanation of anxious states in healthy individuals and anxiety disorders alike (Grupe and Nitschke, 2013). Accordingly, our findings that anxiety leads to changes in informational and environmental uncertainty could prove relevant for understanding the alterations in decision-making and learning observed in anxiety disorders (Bishop and Gagne, 2018; Browning et al., 2015; de Visser et al., 2010; Huang et al., 2017; Miu et al., 2008).

Our approach is not the first in proposing a role of uncertainty estimates in cognitive biases in anxiety. A recent account of affective disorders suggested that difficulties with uncertainty estimation underlie some of the psychiatric symptoms in these populations (Pulcu and Browning, 2019). This work distinguished between different types of uncertainty, corresponding to irreducible, informational, and environmental uncertainty as described here, and assigned a particular relevance of environmental (“unexpected”) uncertainty in explaining anxiety. In fact, evidence from computational studies converges in linking trait anxiety with difficulties in learning in unstable or volatile environments (Browning et al., 2015; Huang et al., 2017). As shown by Browning et al. (2015), an inability to adapt to changes in a task structure can be measured by comparing a single volatile block to a single stable block. Alternatively, suboptimal learning in anxiety can be captured by focusing on volatile environments alone, in which the probability of reward (or punishment) changes regularly across different blocks (Huang et al., 2017). Here we followed the second approach to investigate reward-based learning in a volatile environment. Critically, we investigate adaptive scaling of learning rates to estimates of environmental uncertainty on a trial-by-trial basis by applying a computational model that explicitly incorporates learning about volatility in a hierarchical Bayesian framework. The winning computational model that best explained our behavioural data was the 3-level HGF, where the third level is a mathematical description of volatility estimates and their variance. This model allowed us to assess the effect of state anxiety on an array of relevant computational quantities during task performance. Above and beyond revealing a misestimation of volatility (environmental uncertainty) – as proposed by Pulcu and Browning (2019) – our approach identified biases in uncertainty on various levels that drive suboptimal learning in state anxiety.

First of all, the participants’ estimates of tonic volatility – as captured by the parameter ω_2_ – were significantly reduced in the state anxiety group, which led to significantly reduced learning rates and estimates of informational and environmental uncertainty. Beliefs about the outcome tendency were thus estimated to be more precise during anxiety, such that new and potentially revealing information about the true nature of hidden states had a smaller influence on the belief updates on that level. Critically, an overly precise belief about the outcome tendency might be inappropriate given the fluctuations in the true underlying hidden state. Thus, an aberrant drop in informational uncertainty might lead to biased learning, which here was further characterised by a lower learning rate about stimulus outcomes. This finding was confirmed in a behavioural measure independent of the modelling approach: state anxious individuals exhibited a higher error rate during task performance relative to control participants. Our study thus provides novel and compelling evidence for abnormal precision estimates underlying impoverished learning in healthy individuals going through temporary states of anxiety. Thus, improper precision weighting could be a general mechanism underlying a range of cognitive biases observed in healthy and psychiatric conditions, such as “hysteria” or autism (Edwards et al., 2012; Lawson et al., 2017).

Secondly, we found that state anxiety led to a decrease in the precision of beliefs about environmental volatility. In the HGF update equations, greater uncertainty on the higher level leads to new information having a stronger influence on the update of beliefs concerning volatility (equation 4; see also |Mathys, 2011, 2014), thus rendering this belief more changeable. A greater uncertainty about the world’s current level of stability may underlie an increased tendency to anticipate potential danger. In the context of anxious individuals having negative reactions to uncertainty (Carleton, 2016), our results may reflect how state anxiety increases the likelihood of misinterpreting unstable signals from the environment as threatening – signaling a more volatile world overall – leading to inappropriate value estimates.

In sum, state anxious individuals in our study showed both a decrease in uncertainty and learning rate on the lower level, driven by changes in tonic volatility estimates, ω_2_, and an increase in uncertainty (and thus learning rate) on the higher, volatility estimating level. A possible interpretation of these findings is that rather than learning about (changes of) states themselves, individuals in an anxious state attribute any detected changes in the environment to a potential change in the stability of the world. Lower estimates of tonic volatility (ω_2_) indicate that they expect (or tolerate) fewer changes to the current contingencies governing their environment; instead, when conditions change, they might rapidly infer that the world has gone from a stable to a volatile period due to aberrant uncertainty about the world’s level of stability.

Overall, the computational results confirm our hypothesis that state anxious individuals choose their responses founded on a biased representation of uncertainty over the current belief states – at least when dealing with volatile environments as assessed here. Entertaining overly precise beliefs may represent a strategy to regain a sense of control because uncertainty is experienced as aversive, such as observed in obsessive compulsive disorder (Carleton, 2016) and ritualistic behaviour (Lang et al., 2015). In turn, this emergence of biased estimates could increase the symptoms of anxiety over time through recursive inaccurate assessments of threat from uncertainty, thereby fitting a profile of anxious responses similar to those of anxiety-related disorders (Grupe and Nitschke, 2013; Pulcu and Browning, 2019).

### Hierarchically-related prediction errors modulate trial-by-trial ERP responses

In the control group, both the lower and higher-level pwPE trajectories modulated the trial-by-trial ERP responses. Lower level pwPEs about the reward outcomes, updating beliefs about reward tendency and higher-level pwPEs about reward tendencies, updating volatility estimates, were primarily represented across frontocentral regions, with an earlier latency for ε_3_ (387 ms) than ε_2_ (453 ms). These results align with previous studies combining EEG analyses with the HGF, which revealed that multiple, hierarchically-related pwPEs are computed while learning in volatile environments; and these are represented across different brain regions specific to the task demands (Diaconescu et al., 2017a; Stefanics et al., 2018; Weber et al., 2019).

Importantly, however, this prior work emphasised a specific temporal hierarchy governing the neural representation of the pwPEs across different levels, with lower pwPEs emerging earlier in time relative to higher-level pwPEs. The reasoning behind this emphasis was that in the one-step update equations of the HGF, the lower-level PE needs to be computed first, because the higher-level PE depends on the belief update on the lower level. Here, however, we find that the peak of activation by the higher-level PE precedes that of the lower-level PE. Further work is needed to clarify the implications of this result; however, it may be helpful to consider the nature of the model (HGF) versus the signal (EEG) in this context. The HGF update equations quantify the (total) change in beliefs – both in the mean and the uncertainty or precision – on different levels, in response to an observation. However, while these equations calculate the new posterior in one step, the brain operates in continuous time, where the constant message passing among hierarchically organised regions results in oscillatory signals (Bogacz, 2017; Friston, 2005) which we measure as evoked responses in the EEG. It thus seems likely that while the final value of the posterior belief has to be reached by the end of the ERP, when the evoked oscillation stabilises at a new level, the temporal dynamics leading up to this might be more complex than a linear sequence of updates. Future work on this might profit from formulating an explicit response model which directly links belief states in the HGF to observable EEG responses.

More generally however, the evidence for distinct neural representations of different types of pwPEs in the control group lends support to current predictive coding proposals. These view the brain as a Bayesian observer, estimating beliefs about hidden states in the environment through implementing a hierarchical generative model of the incoming sensory data (de Lange et al., 2018; Doya et al., 2007; Friston, 2010; Rao and Ballard, 1999). In this framework, superficial pyramidal cells encode PEs weighted by precision, and these are also the signals that are thought to dominate the EEG (Friston and Kiebel, 2009). This motivated us to assess the representation of pwPEs in brain responses, an approach followed by some of the previous fMRI and EEG studies (Iglesias et al., 2013; Stefanics et al., 2018; Weber et al., 2019). Other model-based studies of trial-wise ERP responses like the P300 assessed alternative Bayesian inference parameters, such as precision or Bayesian surprise (Kolossa et al., 2015; Mars et al., 2008; Ostwald et al., 2012). The more anterior P3a component around 340 ms was identified as an index of belief updating, whereas the later P3b waveform of posterior topography was found to represent Bayesian surprise (Kolossa et al., 2015). Despite these computational approaches to the P300 not being directly comparable to our pwPE results, they do resemble the timeline and topography of the results from the standard lose minus win ERP analysis conducted here in the Cont and StA groups separately, showing the expected anterior to posterior topographic shift in the P300 component from classical model-free ERP studies (Hajcak et al., 2007, 2005; Polich, 2007; Wu and Zhou, 2009).

Under state anxiety, a neural representation of pwPEs emerged exclusively for volatility updates. Due to the smaller variance of ε_2_ in the anxiety group, it is likely that this regressor was less potent to explain the variance in the EEG signals, making it potentially harder to detect in the brain responses in this group. Possibly, our study was underpowered to detect these weaker effects on the anxiety-related ERP waveforms. The sample size was based on previous research in trait anxiety combining behavioural and computational analysis (Browning et al., 2015), and future work should carry out a sample size estimation specifically targeting the GLM EEG analysis to validate these results. In summary, in state anxiety, belief updating during reward-based learning was mainly driven by updates in volatility, with a corresponding neural representation in parietal electrode regions.

The latency of the trial-wise ERP changes to ε_3_ in anxiety, 359 ms, resembled the main ε_3_ effect in control participants, yet the distribution was shifted towards parietal regions. An important limitation in our study is that we cannot directly statistically compare the effects of the pwPEs on EEG responses between the two groups, because the subject-specific trial-wise regressors used in the GLMs already differ between the groups, due to the effect of state anxiety on belief updating as discussed above. Accordingly, here, we only speculate about explanations for the different pattern of activations observed under state anxiety.

Activity in the right dorsal ACC has been shown to express changes in ε_3_ in a combined fMRI-EEG study (Diaconescu et al., 2017a), while earlier fMRI work linked the ACC, together with the DLPFC, insula and dopaminergic ventral tegmental area / substantia nigra (VTA/SN) to lower-level pwPEs instead; and ε_3_ to the cholinergic basal forebrain (Iglesias et al., 2013). Interpretations with regard to neuroanatomical regions are limited in our EEG study as it provided exclusively sensor-level results. Despite this limitation, potential contributions by the ACC and DLPFC regions to ε_2_ and ε_3_ responses would be compatible with the anterior distribution of pwPEs observed in control participants. Intriguingly, state anxiety has been shown to deactivate the DLPFC (and ventrolateral PFC) and ACC during cognitive control tasks that crucially depend on these areas (Bishop, 2009, 2007; Bishop et al., 2004). Thus, one hypothesis that could be tested in future combined fMRI-EEG studies is whether state anxiety disengages these brain regions during reward-based learning, undermining their proper contribution to tracking pwPE about the reward tendency and volatility. Parietal regions could be playing a compensatory role in the StA group, at least to track pwPE about volatility, whereas any available resources in prefrontal and ACC regions may have been allocated to an *earlier* processing of win and lose outcomes, as shown in the anterior topography of the ERP modulations at 293 ms associated with the outcome regressor. Of particular interest, decreased DLPFC activity also characterises elevated trait anxiety levels, with detrimental consequences for performance and attentional control (Bishop, 2009). And portions of the cingulate cortex and prefrontal cortex are part of the central network underlying anxiety disturbances (Grupe and Nitschke, 2013). Thus, an additional interesting question for future studies would be to assess the role that these brain regions play in the modulation of hierarchically-related pwPEs that may lead to the computational biases described in trait anxiety (Browning et al., 2015; Huang et al., 2017).

## 5. Conclusion and outlook

This study is the first to provide a mechanistic understanding of how temporary anxious states impair reward-based learning in volatile environments. The results thus have implications for understanding cognitive biases and impaired learning in healthy individuals exposed to upcoming uncertain threats, but could also generalise to clinical settings. One important direction for future research will be to determine whether the distributed network of brain regions and neurotransmitter systems linked to anxiety-disorders and trait anxiety interact with the neural representation of hierarchical pwPEs, thereby accounting for the misestimation of both precision and uncertainty, impairing learning.

## Author contributions

M.H.R. and T.P.H. designed the experiment, T.P.H collected the data, M.H.R. and T.P.H. analysed the data, M.H.R., L.A.W. and T.P.H. wrote code for data analysis, M.H.R., L.A.W. and T.P.H. wrote the manuscript, J.D.F. edited the manuscript.

## Acknowledgements

Funding: The study was supported by Goldsmiths University of London, funded by the Economic and Social Research Council (ESRC) and the South East Network for Social Sciences (SeNSS) through grant ES/P00072X/1, and the British Academy, through grant SG161006.

---

**Supplementary Figure 1.**
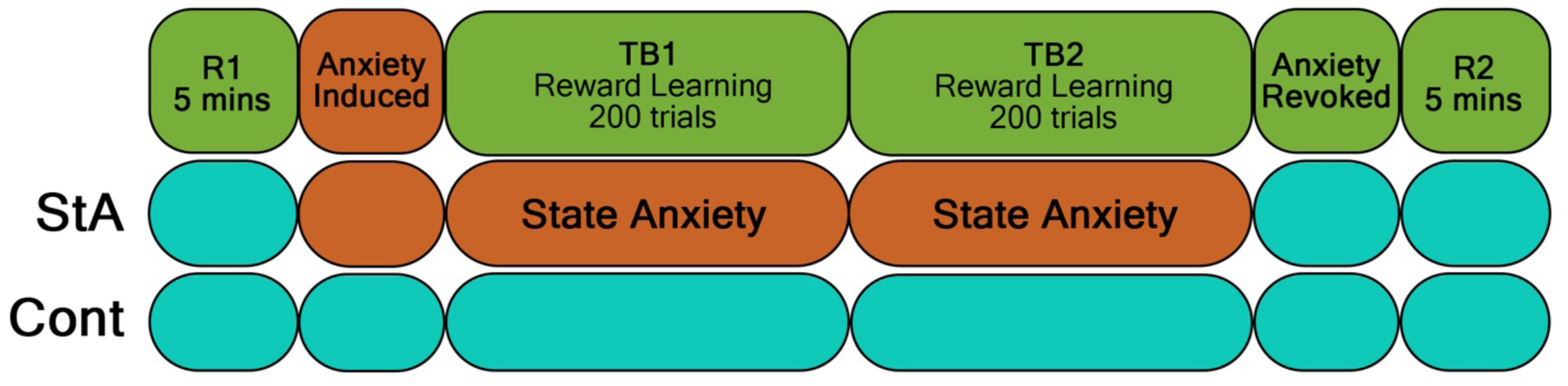
Scheme of the experimental block structure. The experiment was split into four blocks: the first resting-state block (R1: baseline), task block 1 (TB1), task block 2 (TB2), and the final resting state block (R2). R1 comprised of resting-state EEG and ECG recording for 5 minutes in both the state anxiety (StA) and control (Cont) groups. Prior to TB1, the StA group were informed of the experimental manipulation of public speaking about an unknown artwork in front of 3 experts, to be carried out after the computer learning task (TB1, TB2) had finished. This aimed to induce anxiety during TB1 and TB2 for the StA group, as indicated by hatched lines. The Cont group were told to expect to describe the same artwork to themselves for an identical period of time. Both groups undertook the reward-learning task across two blocks (TB1, TB2). After completing TB2, the StA group were informed they would not need to speak publicly about the artwork, and that they would, in an identical fashion to the Cont group, only need to present the artwork to themselves. When finished with this self-presentation, the final resting state block of ECG and EEG was recorded (R2).

**Supplementary Figure 2.**
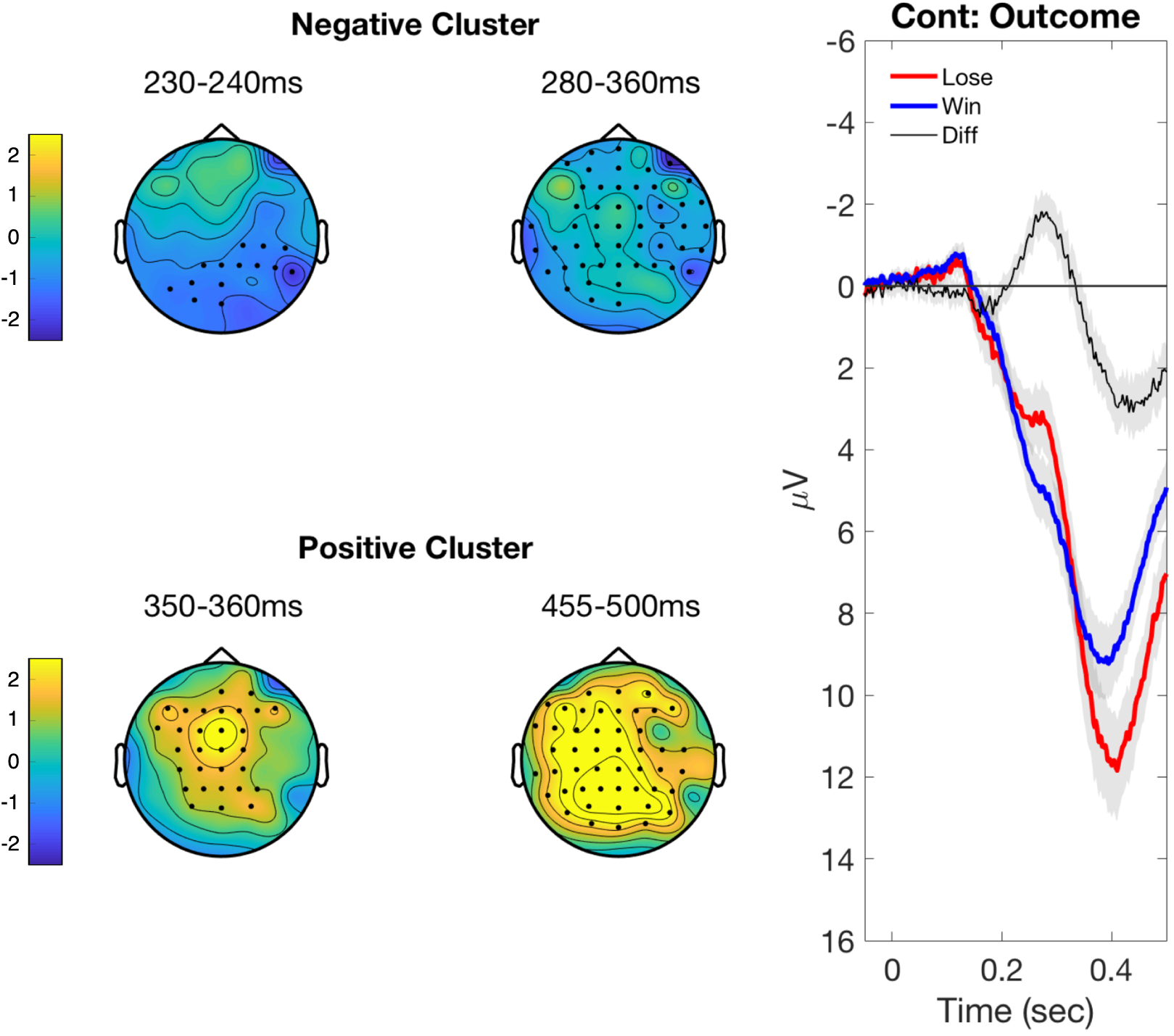
**Results of the ERP response comparison between outcome in the control group: wins and losses. Left panels**. Cluster-based random permutation analysis of ERP responses was carried out in the total population of the control group (Cont, N= 21) to assess the effect of the outcome (win, lose). Maps given for each cluster show the scalp topography of the significant cluster ERP differences between outcomes (win, lose) across an earlier (leftmost) time window and later (rightmost) time window. We give these two time frames to demonstrate how the significant clusters propagate. Black dots on the topographical maps indicate electrodes pertaining to a significant cluster (*P* < 0.025, two-tailed test). **Right panel**. Grand-mean ERP waveforms of the two outcomes (lose, red; win, blue) and the difference (lose minus win, black) are presented from all electrodes between −0.05 and 0.5 seconds, with SEM given as grey shaded areas. Significant clusters are denoted by black bars on the x-axis.

**Supplementary Figure 3.**
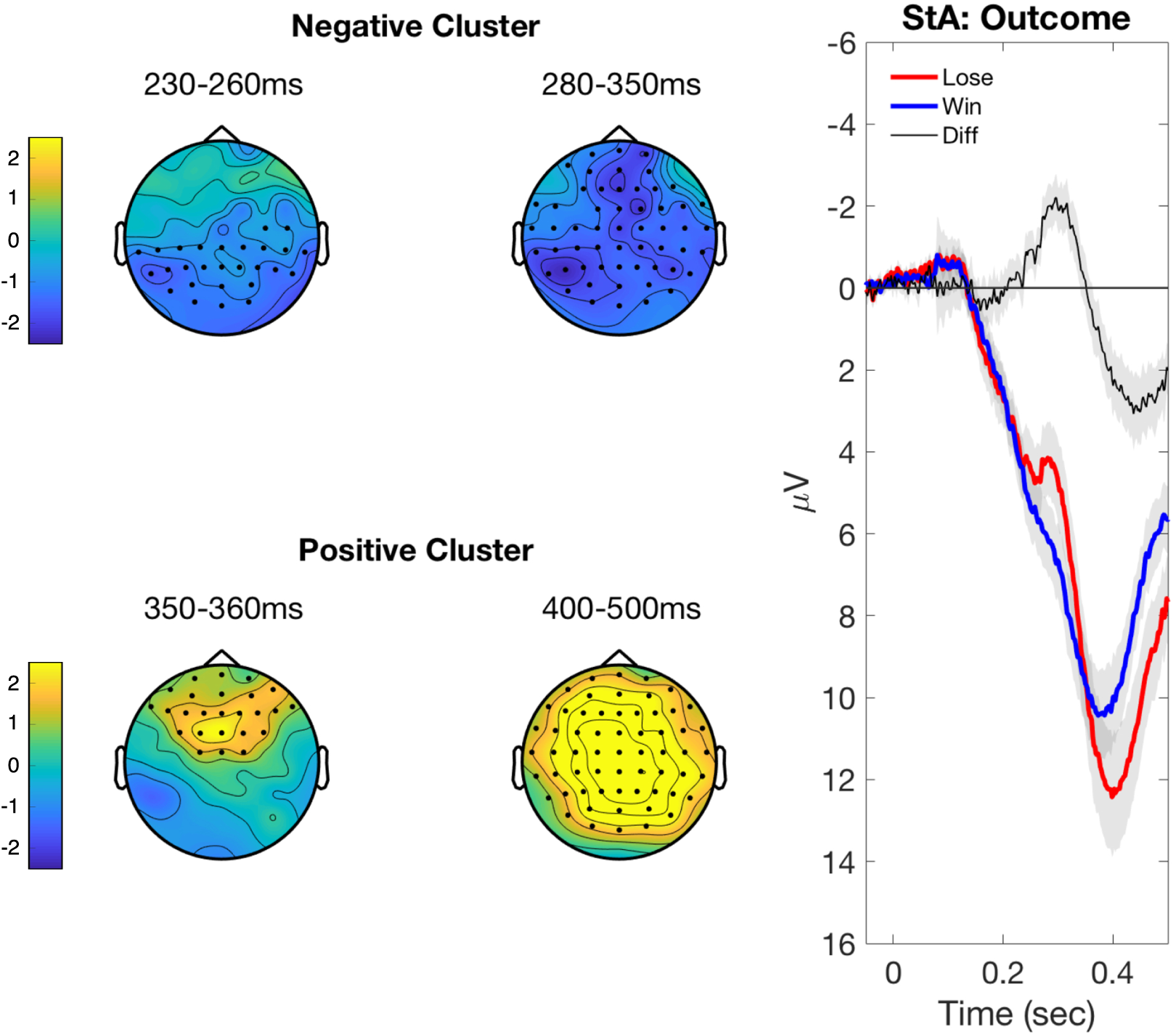
**Results of the ERP response comparison between outcome in the state anxiety group: wins and losses. Left panels**. Cluster-based random permutation analysis of ERP responses in the state anxiety group (StA, N= 21) to assess the effect of the outcome (win, lose). Maps given for each cluster show the scalp topography of the significant cluster ERP differences between outcomes (win, lose) across an earlier (leftmost) time window and later (rightmost) time window. We present this to show the spread of the clusters. Black dots on the topographical maps indicate electrodes referring to a significant cluster (*P* < 0.025, two-tailed test). **Right panel**. Grand-mean ERP waveforms of the two outcomes (lose, red; win, blue) and the difference (lose minus win, black) are presented from all electrodes between −0.05 and 0.5 seconds, with SEM given as grey shaded areas. Significant clusters are denoted by black bars on the x-axis.

**Supplementary Figure 4.**
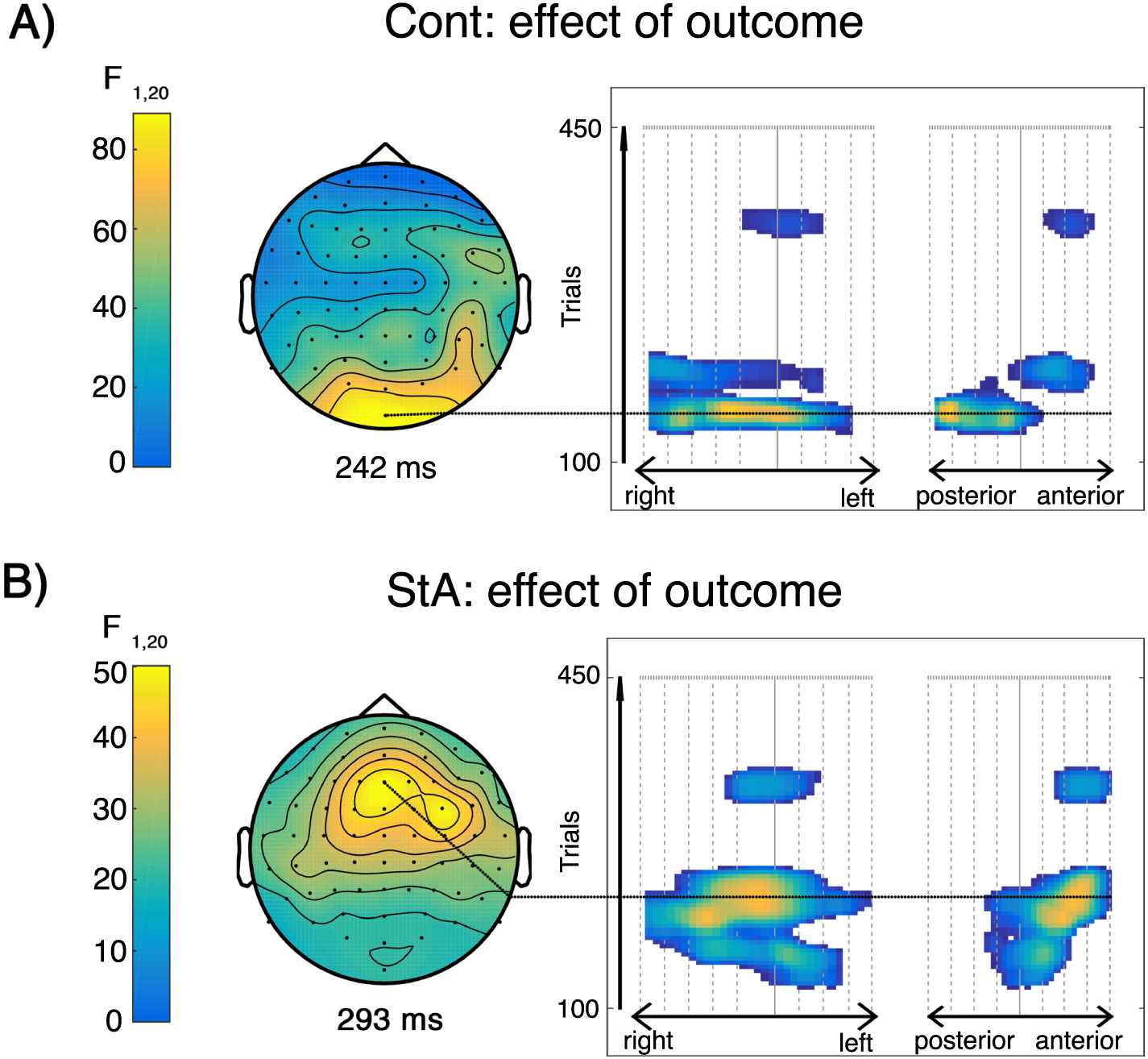
**Effect of the outcome regressor on trial-wise ERPs**. **A)** The effect of trial outcome on EEG brain activity in the control group. Using trial outcomes (win, loss) as a regressor, a significant representation in ERP responses were shown between 226 ms to 259 ms over parieto-occipital regions, with the peak effect occurring at 242 ms (*P_FWE_* < 0.0001, with a cluster-defining threshold of P < 0.001). An additional effect included a later cluster in frontal locations between 404-424 ms (*P_FWE_* = 0.004). **B)** The effect of trial outcome on EEG brain activity in the state anxiety group. In this group, the regressor corresponding to the trial outcomes led to a significant modulation of the ERP waveform between 254 ms and 322 ms at frontal and fronto-central channels, with the peak effect occurring at 293 ms (*P_FWE_* = 0, with a cluster-defining threshold of *P* < 0.001). In addition, a later significant cluster in the same frontal-central channels was found between 388 and 416 ms, with a maximum peak at 402 ms (*P_FWE_* = 0.003).

## References

Bach, D.R., Hulme, O., Penny, W.D., Dolan, R.J., 2011. The known unknowns: neural representation of second-order uncertainty, and ambiguity. J. Neurosci. 31, 4811–4820.

Barnes, L.L.B., Harp, D., Jung, W.S., 2002. Reliability Generalization of Scores on the Spielberger State-Trait Anxiety Inventory. Educ. Psychol. Meas. 62, 603–618.

Basso, D., Chiarandini, M., Salmaso, L., 2007. Synchronized permutation tests in replicated I×J designs. J. Stat. Plan. Inference 137, 2564–2578.

Bastos, A.M., Usrey, W.M., Adams, R.A., Mangun, G.R., Fries, P., Friston, K.J., 2012. Canonical microcircuits for predictive coding. Neuron 76, 695–711.

Behrens, T.E.J., Woolrich, M.W., Walton, M.E., Rushworth, M.F.S., 2007. Learning the value of information in an uncertain world. Nat. Neurosci. 10, 1214–1221.

Benjamini, Y., Krieger, A.M., Yekutieli, D., 2006. Adaptive linear step-up procedures that control the false discovery rate. Biometrika 93, 491–507.

Bishop, S.J., 2009. Trait anxiety and impoverished prefrontal control of attention. Nat. Neurosci. 12, 92–98.

Bishop, S.J., 2008. Neural mechanisms underlying selective attention to threat. Ann. N. Y. Acad. Sci. 1129, 141–152.

Bishop, S.J., 2007. Neurocognitive mechanisms of anxiety: an integrative account. Trends Cogn. Sci. 11, 307–316.

Bishop, S.J., Duncan, J., Lawrence, A.D., 2004. State anxiety modulation of the amygdala response to unattended threat-related stimuli. J. Neurosci. 24, 10364–10368.

Bishop, S.J., Gagne, C., 2018. Anxiety, Depression, and Decision Making: A Computational Perspective. Annu. Rev. Neurosci. 41, 371–388.

Bland, A.R., Schaefer, A., 2012. Different varieties of uncertainty in human decision-making. Front. Neurosci. 6, 85.

Bogacz, R., 2017. A tutorial on the free-energy framework for modelling perception and learning. J. Math. Psychol. 76, 198–211.

Browning, M., Behrens, T.E., Jocham, G., O’Reilly, J.X., Bishop, S.J., 2015. Anxious individuals have difficulty learning the causal statistics of aversive environments. Nat. Neurosci. 18, 590–596.

Carleton, R.N., 2016. Into the unknown: A review and synthesis of contemporary models involving uncertainty. J. Anxiety Disord. 39, 30–43.

Chalmers, J.A., Quintana, D.S., Abbott, M.J.-A., Kemp, A.H., 2014. Anxiety Disorders are Associated with Reduced Heart Rate Variability: A Meta-Analysis. Front. Psychiatry 5, 80.

Chew, B., Hauser, T.U., Papoutsi, M., Magerkurth, J., Dolan, R.J., Rutledge, R.B., 2019. Endogenous fluctuations in the dopaminergic midbrain drive behavioral choice variability. Proc. Natl. Acad. Sci. U. S. A. https://doi.org/10.1073/pnas.1900872116

Cisler, J.M., Koster, E.H.W., 2010. Mechanisms of attentional biases towards threat in anxiety disorders: An integrative review. Clin. Psychol. Rev. 30, 203–216.

de Berker, A.O., Rutledge, R.B., Mathys, C., Marshall, L., Cross, G.F., Dolan, R.J., Bestmann, S., 2016. Computations of uncertainty mediate acute stress responses in humans. Nat. Commun. 7, 10996.

de Lange, F.P., Heilbron, M., Kok, P., 2018. How Do Expectations Shape Perception? Trends Cogn. Sci. 22, 764–779.

Delorme, A., Makeig, S., 2004. EEGLAB: an open source toolbox for analysis of single-trial EEG dynamics including independent component analysis. J. Neurosci. Methods 134, 9–21.

de Visser, L., van der Knaap, L.J., van de Loo, A.J.A.E., van der Weerd, C.M.M., Ohl, F., van den Bos, R., 2010. Trait anxiety affects decision-making differently in healthy men and women: towards gender-specific endophenotypes of anxiety. Neuropsychologia 48, 1598–1606.

Diaconescu, A.O., Litvak, V., Mathys, C., Kasper, L., Friston, K.J., Stephan, K.E., 2017a. A computational hierarchy in human cortex. arXiv [q-bio.NC].

Diaconescu, A.O., Mathys, C., Weber, L.A.E., Daunizeau, J., Kasper, L., Lomakina, E.I., Fehr, E., Stephan, K.E., 2014. Inferring on the intentions of others by hierarchical Bayesian learning. PLoS Comput. Biol. 10, e1003810.

Diaconescu, A.O., Mathys, C., Weber, L.A.E., Kasper, L., Mauer, J., Stephan, K.E., 2017b. Hierarchical prediction errors in midbrain and septum during social learning. Soc. Cogn. Affect. Neurosci. 12, 618–634.

Doya, K., Ishii, S., Pouget, A., Rao, R.P.N., 2007. Bayesian Brain: Probabilistic Approaches to Neural Coding. MIT Press.

Edwards, M.J., Adams, R.A., Brown, H., Pareés, I., Friston, K.J., 2012. A Bayesian account of “hysteria.” Brain 135, 3495–3512.

Feldman, H., Friston, K.J., 2010. Attention, uncertainty, and free-energy. Front. Hum. Neurosci. 4, 215.

Feldman, P.J., Cohen, S., Hamrick, N., Lepore, S.J., 2004. Psychological stress, appraisal, emotion and Cardiovascular response in a public speaking task. Psychol. Health 19, 353– 368.

Flandin, G., Friston, K.J., 2019. Analysis of family-wise error rates in statistical parametric mapping using random field theory. Hum. Brain Mapp. 40, 2052–2054.

Friedman, B.H., 2007. An autonomic flexibility–neurovisceral integration model of anxiety and cardiac vagal tone. Biol. Psychol. 74, 185–199.

Friedman, B.H., Thayer, J.F., 1998. Autonomic balance revisited: panic anxiety and heart rate variability. J. Psychosom. Res. 44, 133–151.

Friston, K., 2005. A theory of cortical responses. Philos. Trans. R. Soc. Lond. B Biol. Sci. 360, 815–836.

Friston, K., 2010. The free-energy principle: a unified brain theory? Nat. Rev. Neurosci. 11, 127–138.

Friston, K., Brown, H.R., Siemerkus, J., Stephan, K.E., 2016. The dysconnection hypothesis. Schizophr. Res. 176, 83–94.

Friston, K., Kiebel, S., 2009. Predictive coding under the free-energy principle. Philos. Trans. R. Soc. Lond. B Biol. Sci. 364, 1211–1221.

Friston, K., Schwartenbeck, P., Fitzgerald, T., Moutoussis, M., Behrens, T., Dolan, R.J., 2013. The anatomy of choice: active inference and agency. Front. Hum. Neurosci. 7, 598.

Gehring, W.J., Willoughby, A.R., 2004. Are all medial frontal negativities created equal? Toward a richer empirical basis for theories of action monitoring. Errors, conflicts, and the brain. Current opinions on performance monitoring 14, 20.

Gorman, J.M., Sloan, R.P., 2000. Heart rate variability in depressive and anxiety disorders. Am. Heart J. 140, 77–83.

Grissom, R.J., Kim, J.J., 2012. Effect sizes for research: Univariate and multivariate applications. Routledge.

Grupe, D.W., Nitschke, J.B., 2013. Uncertainty and anticipation in anxiety: an integrated neurobiological and psychological perspective. Nat. Rev. Neurosci. 14, 488–501.

Hajcak, G., Holroyd, C.B., Moser, J.S., Simons, R.F., 2005. Brain potentials associated with expected and unexpected good and bad outcomes. Psychophysiology 42, 161–170.

Hajcak, G., Moser, J.S., Holroyd, C.B., Simons, R.F., 2007. It’s worse than you thought: the feedback negativity and violations of reward prediction in gambling tasks. Psychophysiology 44, 905–912.

Holroyd, C.B., Coles, M.G.H., 2008. Dorsal anterior cingulate cortex integrates reinforcement history to guide voluntary behavior. Cortex 44, 548–559.

Holroyd, C.B., Krigolson, O.E., 2007. Reward prediction error signals associated with a modified time estimation task. Psychophysiology 44, 913–917.

Holroyd, C.B., Nieuwenhuis, S., Yeung, N., Cohen, J.D., 2003. Errors in reward prediction are reflected in the event-related brain potential. Neuroreport 14, 2481–2484.

Huang, H., Thompson, W., Paulus, M.P., 2017. Computational Dysfunctions in Anxiety: Failure to Differentiate Signal From Noise. Biol. Psychiatry 82, 440–446.

Iglesias, S., Mathys, C., Brodersen, K.H., Kasper, L., Piccirelli, M., den Ouden, H.E.M., Stephan, K.E., 2013. Hierarchical prediction errors in midbrain and basal forebrain during sensory learning. Neuron 80, 519–530.

Jepma, M., Murphy, P.R., Nassar, M.R., Rangel-Gomez, M., Meeter, M., Nieuwenhuis, S., 2016. Catecholaminergic Regulation of Learning Rate in a Dynamic Environment. PLoS Comput. Biol. 12, e1005171.

Kawachi, I., Sparrow, D., Vokonas, P.S., Weiss, S.T., 1995. Decreased heart rate variability in men with phobic anxiety (data from the Normative Aging Study). Am. J. Cardiol. 75, 882– 885.

Kiebel, S.J., Friston, K.J., 2004a. Statistical parametric mapping for event-related potentials: I. Generic considerations. Neuroimage 22, 492–502.

Kiebel, S.J., Friston, K.J., 2004b. Statistical parametric mapping for event-related potentials (II): a hierarchical temporal model. Neuroimage 22, 503–520.

Kilner, J.M., Friston, K.J., 2010. Topological inference for EEG and MEG. Ann. Appl. Stat. 4, 1272–1290.

Kok, P., de Lange, F.P., 2015. Predictive Coding in Sensory Cortex, in: Forstmann, B.U., Wagenmakers, E.-J. (Eds.), An Introduction to Model-Based Cognitive Neuroscience. Springer New York, New York, NY, pp. 221–244.

Kolossa, A., Kopp, B., Fingscheidt, T., 2015. A computational analysis of the neural bases of Bayesian inference. Neuroimage 106, 222–237.

Kriegeskorte, N., Simmons, W.K., Bellgowan, P.S.F., Baker, C.I., 2009. Circular analysis in systems neuroscience: the dangers of double dipping. Nat. Neurosci. 12, 535–540.

Lang, M., Krátký, J., Shaver, J.H., Jerotijević, D., Xygalatas, D., 2015. Effects of Anxiety on Spontaneous Ritualized Behavior. Curr. Biol. 25, 1892–1897.

Lawson, R.P., Mathys, C., Rees, G., 2017. Adults with autism overestimate the volatility of the sensory environment. Nat. Neurosci. 20, 1293–1299.

Lawson, R.P., Rees, G., Friston, K.J., 2014. An aberrant precision account of autism. Front. Hum. Neurosci. 8, 302.

Litvak, V., Mattout, J., Kiebel, S., Phillips, C., Henson, R., Kilner, J., Barnes, G., Oostenveld, R., Daunizeau, J., Flandin, G., Penny, W., Friston, K., 2011. EEG and MEG data analysis in SPM8. Comput. Intell. Neurosci. 2011, 852961.

Lorberbaum, J.P., Kose, S., Johnson, M.R., Arana, G.W., Sullivan, L.K., Hamner, M.B., Ballenger, J.C., Lydiard, R.B., Brodrick, P.S., Bohning, D.E., George, M.S., 2004. Neural correlates of speech anticipatory anxiety in generalized social phobia. Neuroreport 15, 2701–2705.

Maris, E., Oostenveld, R., 2007. Nonparametric statistical testing of EEG- and MEG-data. J. Neurosci. Methods 164, 177–190.

Marshall, L., Mathys, C., Ruge, D., de Berker, A.O., Dayan, P., Stephan, K.E., Bestmann, S., 2016. Pharmacological Fingerprints of Contextual Uncertainty. PLoS Biol. 14, e1002575.

Mars, R.B., Debener, S., Gladwin, T.E., Harrison, L.M., Haggard, P., Rothwell, J.C., Bestmann, S., 2008. Trial-by-trial fluctuations in the event-related electroencephalogram reflect dynamic changes in the degree of surprise. J. Neurosci. 28, 12539–12545.

Mathys, C., Daunizeau, J., Friston, K.J., Stephan, K.E., 2011. A bayesian foundation for individual learning under uncertainty. Front. Hum. Neurosci. 5, 39.

Mathys, C., Lomakina, E.I., Daunizeau, J., Iglesias, S., Brodersen, K.H., Friston, K.J., Stephan, K.E., 2014. Uncertainty in perception and the Hierarchical Gaussian Filter. Front. Hum. Neurosci. 8, 825.

Matsumoto, M., Matsumoto, K., Abe, H., Tanaka, K., 2007. Medial prefrontal cell activity signaling prediction errors of action values. Nat. Neurosci. 10, 647–656.

Miu, A.C., Heilman, R.M., Houser, D., 2008. Anxiety impairs decision-making: psychophysiological evidence from an Iowa Gambling Task. Biol. Psychol. 77, 353–358.

Montague, P.R., Hyman, S.E., Cohen, J.D., 2004. Computational roles for dopamine in behavioural control. Nature 431, 760–767.

Moody, G.B., Mark, R.G., 1982. Development and evaluation of a 2-lead ECG analysis program. Comput. Cardiol. 9, 39–44.

Moran, R.J., Campo, P., Symmonds, M., Stephan, K.E., Dolan, R.J., Friston, K.J., 2013. Free energy, precision and learning: the role of cholinergic neuromodulation. J. Neurosci. 33, 8227–8236.

Morris, G., Nevet, A., Arkadir, D., Vaadia, E., Bergman, H., 2006. Midbrain dopamine neurons encode decisions for future action. Nat. Neurosci. 9, 1057–1063.

Nieuwenhuis, S., Holroyd, C.B., Mol, N., Coles, M.G.H., 2004. Reinforcement-related brain potentials from medial frontal cortex: origins and functional significance. Neurosci. Biobehav. Rev. 28, 441–448.

Oostenveld, R., Fries, P., Maris, E., Schoffelen, J.-M., 2011. FieldTrip: Open source software for advanced analysis of MEG, EEG, and invasive electrophysiological data. Comput. Intell. Neurosci. 2011, 156869.

Ostwald, D., Spitzer, B., Guggenmos, M., Schmidt, T.T., Kiebel, S.J., Blankenburg, F., 2012. Evidence for neural encoding of Bayesian surprise in human somatosensation. Neuroimage 62, 177–188.

Parr, T., Rees, G., Friston, K.J., 2018. Computational Neuropsychology and Bayesian Inference. Front. Hum. Neurosci. 12, 61.

Paulus, M.P., Stein, M.B., 2010. Interoception in anxiety and depression. Brain Struct. Funct. 214, 451–463.

Paulus, M.P., Yu, A.J., 2012. Emotion and decision-making: affect-driven belief systems in anxiety and depression. Trends Cogn. Sci. 16, 476–483.

Penny, W.D., Friston, K.J., Ashburner, J.T., Kiebel, S.J., Nichols, T.E., 2011. Statistical Parametric Mapping: The Analysis of Functional Brain Images. Elsevier.

Penny, W., Holmes, A., 2007. Random effects analysis. Statistical parametric mapping: The analysis of functional brain images 156–165.

Polich, J., 2007. Updating P300: An integrative theory of P3a and P3b. Clin. Neurophysiol. 118, 2128–2148.

Pulcu, E., Browning, M., 2019. The Misestimation of Uncertainty in Affective Disorders. Trends Cogn. Sci. https://doi.org/10.1016/j.tics.2019.07.007

Quintana, D.S., Alvares, G.A., Heathers, J.A.J., 2016. Guidelines for Reporting Articles on Psychiatry and Heart rate variability (GRAPH): recommendations to advance research communication. Transl. Psychiatry 6, e803.

Rao, R.P., Ballard, D.H., 1999. Predictive coding in the visual cortex: a functional interpretation of some extra-classical receptive-field effects. Nat. Neurosci. 2, 79–87.

Rescorla, R.A., Wagner, A.R., 1972. A theory of Pavlovian conditioning: Variations in the effectiveness of reinforcement and nonreinforcement. Classical conditioning II: Current research and theory 2, 64–99.

Ruscio, J., Mullen, T., 2012. Confidence Intervals for the Probability of Superiority Effect Size Measure and the Area Under a Receiver Operating Characteristic Curve. Multivariate Behav. Res. 47, 201–223.

Soch, J., Allefeld, C., 2018. MACS--a new SPM toolbox for model assessment, comparison and selection. J. Neurosci. Methods.

Soch, J., Haynes, J.-D., Allefeld, C., 2016. How to avoid mismodelling in GLM-based fMRI data analysis: cross-validated Bayesian model selection. Neuroimage 141, 469–489.

Spielberger, C.D., 1983. Manual for the State-Trait Anxiety Inventory STAI (form Y)(“ self-evaluation questionnaire”).

Spielberger, C.D., Gorsuch, R.L., Lushene, R.E., 1970. Manual for the state-trait anxiety inventory.

Stefanics, G., Heinzle, J., Horváth, A.A., Stephan, K.E., 2018. Visual Mismatch and Predictive Coding: A Computational Single-Trial ERP Study. J. Neurosci. 38, 4020–4030.

Stephan, K.E., Penny, W.D., Daunizeau, J., Moran, R.J., Friston, K.J., 2009. Bayesian model selection for group studies. Neuroimage 46, 1004–1017.

Sutton, R.S., 1992. Gain adaptation beats least squares, in: Proceedings of the 7th Yale Workshop on Adaptive and Learning Systems.

Taylor, D.J., Lichstein, K.L., Durrence, H.H., Reidel, B.W., Bush, A.J., 2005. Epidemiology of insomnia, depression, and anxiety. Sleep 28, 1457–1464.

Weber, L., Diaconescu, A.O., Mathys, C., Schmidt, A., Kometer, M., Vollenweider, F., Stephan, K.E., 2019. Ketamine Affects Prediction Errors About Statistical Regularities: A Computational Single-Trial Analysis of the Mismatch Negativity. bioRxiv. https://doi.org/10.1101/528372

Wilson, F.N., Macleod, A.G., Barker, P.S., 1931. The potential variations produced by the heart beat at the apices of Einthoven’s triangle. Am. Heart J. 7, 207–211.

Worsley, K.J., Marrett, S., Neelin, P., Vandal, A.C., Friston, K.J., Evans, A.C., 1996. A unified statistical approach for determining significant signals in images of cerebral activation. Hum. Brain Mapp. 4, 58–73.

Wu, Y., Zhou, X., 2009. The P300 and reward valence, magnitude, and expectancy in outcome evaluation. Brain Res. 1286, 114–122.

Yeung, N., Holroyd, C.B., Cohen, J.D., 2005. ERP correlates of feedback and reward processing in the presence and absence of response choice. Cereb. Cortex 15, 535–544.

Yu, A.J., Dayan, P., 2005. Uncertainty, neuromodulation, and attention. Neuron 46, 681–692.

Zarr, N., Brown, J.W., 2016. Hierarchical error representation in medial prefrontal cortex. Neuroimage 124, 238–247.

Worsley, K.J., Marrett, S., Neelin, P., Vandal, A.C., Friston, K.J., Evans, A.C., 1996. A unified statistical approach for determining significant signals in images of cerebral activation. Hum. Brain Mapp. 4, 58–73. https://doi.org/3.0.CO;2-O">10.1002/(SICI)1097-0193(1996)4:1&lt;58::AID-HBM4>3.0.CO;2-O

